# DNN-DTIs: improved drug-target interactions prediction using XGBoost feature selection and deep neural network

**DOI:** 10.1101/2020.08.11.247437

**Authors:** Cheng Chen, Han Shi, Yu Han, Zhiwen Jiang, Xuefeng Cui, Bin Yu

**Affiliations:** College of Mathematics and Physics, Qingdao University of Science and Technology, Qingdao 266061, China; Artificial Intelligence and Biomedical Big Data Research Center, Qingdao University of Science and Technology, Qingdao 266061, China; Key Laboratory of Synthetic Biology, CAS Center for Excellence in Molecular Plant Sciences, Chinese Academy of Sciences, Shanghai, 200032, China; School of Computer Science and Technology, Shandong University, Qingdao 266237, China; School of Life Sciences, University of Science and Technology of China, Hefei 230027, China

**Keywords:** Drug-target interactions, Multi-information fusion, XGBoost, Deep neural network

## Abstract

Research, analysis, and prediction of drug-target interactions (DTIs) play an important role in understanding drug mechanisms, drug repositioning and design. Machine learning (ML)-based methods for DTIs prediction can mitigate the shortcomings of time-consuming and labor-intensive experimental approaches, providing new ideas and insights for drug design. We propose a novel pipeline for predicting drug-target interactions, called DNN-DTIs. First, the target information is characterized by pseudo-amino acid composition, pseudo position-specific scoring matrix, conjoint triad, composition, transition and distribution, Moreau-Broto autocorrelation, and structure feature. Then, the drug compounds are encoded using substructure fingerprint. Next, we utilize XGBoost to determine nonredundant and important feature subset, then the optimized and balanced sample vectors could be obtained through SMOTE. Finally, a DTIs predictor, DNN-DTIs, is developed based on deep neural network (DNN) via layer-by-layer learning. Experimental results indicate that DNN-DTIs achieves outstanding performance than other predictors with the ACC values of 98.78%, 98.60%, 97.98%, 98.24% and 98.00% on Enzyme, Ion Channels (IC), GPCR, Nuclear Receptors (NR) and Kuang's dataset. Therefore, DNN-DTIs's accurate prediction performance on Network1 and Network2 make it logical choice for contributing to the study of DTIs, especially, the drug repositioning and new usage of old drugs.

## 1. Introduction

The study of drug-target interactions (DTIs) is very helpful for the exploration of the potential beneficial therapeutic effects, the understanding of pharmacology, the prediction of the adverse reactions and drug repositioning. With the development of genomics, proteomics and molecular biology, researchers have built public databases [1–3] through comprehensive knowledge of biology and chemistry, such as Super Target [4], KEGG [5], DrugBank [6], and TTD [7]. However, most of DTIs identified through traditional methods are cumbersome, expensive, and time-consuming. Considering the side effects and toxicity of clinical drugs, only a few drug candidates are approved to enter the market. Therefore, artificial intelligence technology-based prediction methods for DTIs can reduce the waste of manpower and material resources. It also could provide new ideas for repositioning drugs, offering candidate drugs and detecting drug side effects. Especially, it can help to guide to complete experimental task. In recent years, it has become a research focus to predict DTIs with konwledge of drug candidates and specific targets based on computational biology [8,9].

Traditional approaches for predicting DTIs are roughly divided into three categories: ligand-based methods [10], docking methods [11], and chemogenomic methods that comprehensively utilize drug compound and target sequence information [12]. However, the predictions of ligand-based methodology largely depends on prior information of known ligands [13]. The docking methods require the 3D structure of the known target [14]. Both methods have defects, which are time and labor-consuming. The chemogenomic method can be performed according to existing information in public databases, including target sequence-based, structure-based, evolution-based, drug molecular fingerprint-based and biochemistry-based information. In order to facilitate the drug developers and experimenters for in-depth understanding of DTIs, Ezzat et al. [12] divided the chemogenomic methods into neighborhood model, bipartite local models, network diffusion methods, matrix factorization methods, and feature-based classification methods. Bagherian et al. [15] reviewed the machine learning methods for DTIs prediction. Unlike the review methods proposed by Ezzat et al., chemogenomic method can be divided more detailed approaches, including deep learning (DL)-based and hybrid-based methods.

At present, researchers have made a lot of progress and valuable results for DTIs prediction based on machine learning [16]. Yamanishi et al. [17] firstly transformed DTIs predictions into supervised bipartite graph learning methods by integrating chemistry and genomics information. Subsequently, Yamanishi et al. [18] learned pharmacological effects from the similarity of chemical structures, and inferred unknown DTIs from the pharmacological effects and genomic sequence information. Cao et al. [19] proposed an extended structure-activity relationship (SAR) method, which encoded drug molecules with MACCS substructure fingerprints, as well as, encoding target proteins through physicochemical properties. The outputed prediction accuracies through support vector machine are 90.31%, 88.91%, 84.68% and 83.74% on gold standard datasets. Shi et al. [20] enhanced the similarity measure and designed WNN-GIP and KBMF2K by dealing with the missing interaction problem. Ezzat et al. [21] proposed two matrix factorization methods to predict DTIs between “new drugs” and “new targets” by adding edges through intermediate interaction likelihood scores. Hao et al. [22] proposed a simple and effective predictor based on kernel fusion algorithm called RLS-KF, which used regularized least squares integration. The AUPR values of Enzyme, ion channel (IC), GPCR and NR respectively reached 0.915, 0.92.5, 0.853 and 0.909 with 10 times of 10-fold cross-validation. Zhang et al. [23] used the software “PaDEL Descriptor” to describe the drug features, and concatenated with target feature vectors using physicochemical property to construct the raw feature space. Then random projection was employed to generate low-dimensional but effective vectors from original space. Xia et al. [24] used the matrix decomposition method of low-rank weight estimation to predict DTIs, called SPLCMF. They obtained low-dimensional vectors in the drug and target network based on the least squares method, and the soft weight method was employed to reflect the hidden information during training process. They also identified potential DTIs on the four datasets of NR, GPCR, IC and Enzyme. Finally, a DTIs network diagram is ploted to explain its corresponding topology. Li et al. [25] proposed a multi-view low-rank embedding DTIs prediction method, which characterized drug and target information by fusing structural information and chemical information to predict DTIs. Shi et al. [26] proposed a prediction method, LRF-DTIs, using FP2 molecular fingerprint to encode drug and pseudo position-specific scoring matrix to obtain target. They took the optimaized vectors via feature selection and employed SMOTE as input, then outputted the prediction indicators using random forest as classifier. A DTIs identification method proposed by Olayan et al. [27] was called DDR. The nonlinear similarity fusion method was used to extract the similarity matrix information of drugs and targets, and different graph-based feature vectors were extracted from DTIs heterogeneous graph. Finally, random forest was employed to verify the effectiveness of the DDR method.

In the past few years, deep learning has been applied in the field of biomedicine and artificial intelligence at an unprecedented speed, and many DL frameworks have been used to deal with the prediction problem of DTIs [28,29]. A convolutional neural network (CNN)-based method proposed by Öztürk et al. [30], using only sequence information and performing DTIs prediction on Davis and KIBA dataset with the AUPR values of 0.714 and 0.788, respectively. Rayhan et al. [31] proposed the FRnet-DTI, using autoenconder and CNN for feature extraction and classification. The AUC and AUPR values on the gold standard dataset showed that FRnet-DTI could significantly boost the DTIs prediction performance. Lee et al. [32] used a convolutional neural network to extract the local distribution pattern of amino acid sequences, and trained a large-scale DTIs and non-DTIs to build a DL model. This method is superior to the prediction method of DTIs based on deep belief networks. Zeng et al. [33] used cascade deep forest combining with arbitrary-order neighboring algorithms to predict DTIs, thus a network-based computing method called AOPEDF was proposed. The effective feature vectors containing chemical, genomic, phenotypic, and network profiles were inputted into the cascade deep forest. Zhao et al. [34] proposed a new graph convolutional network (GCN)-based method, which extracted feature information of drug-target pairs through GCN. The high-level features were inputted into the deep neural network, and a prediction model based on GCN was constructed.

Although much researches have been done based on ML and DL for drug-target interactions prediction, there is still much room for improvements in the following four areas. First, the number of candidate drugs and candidate targets that do not interact is much larger than the number of drugs and targets that interact, leading to the serious imbalance problem. Second, more and more feature information of drugs and targets apperar, including biological sequence information, network topology information, physicochemical information and structural information. The ways to integrate homogeneous and heterogeneous feature information effectively and comprehensibly are necessary. Third, computational biological-based methods have achieved good prediction results in DTIs prediction. It does not mean that it is effective to predict the interaction between unknown or independent DTIs. It is of great necessity to evaluate the trained model based on DTIs network. Fourth, deep learning, including DNN and CNN, exists many hyperparameters that need to be adjusted. Designing effective deep learning-based pipeline is urgently required for performing efficient and accurate DITs prediction.

Inspired by the above discussion, the DNN-DTIs predictor, based on deep learning and multi-feature fusion, is proposed for DTIs prediction. First, the sequence-based, structure-based and evolution-based target information is derived. Meanwhile, molecular fingerprints is utilized to encode drug compounds. In the prediction field of DTIs, we firstly fuse PseAAC, CT, CTD, NMBroto and secondary structure to elaborate target information. The tree-based XGBoost is employed to determine the optimal feature subset in terms of gain, and then the SMOTE algorithm is used to generate artificial samples for processing the data imbalance problem. It’s worth mentioning that we performs feature selection based on XGBoost to predict DTIs for the first time. Finally, a DNN framework with multilayer hidden layer is constructed. After applying the optimization of different feature extraction method algorithms, feature selection methods and classifier algorithm, the DNN-DTIs model is built up using the optimal parameters. Results on Enzyme, IC, GPCR, NR, and Kuang’s dataset indicates the effectiveness of DNN-DTIs. DNN-DTIs demonstrates the promising drug design validation when it predicts DTIs on Network1 and Network2 successfully, showing the improvement on performance and topological property.

## 2. Materials and methods

### 2.1. Datasets

In this study, seven drug-target datasets are collected, four of which are from the gold standard dataset (GSD) construted by Yamanishi et al. [17]. The datasets are employed to perform parameter optimization, which are divided into four categories: Enzyme, IC, GPCR and NR. The specific distribution of each dataset can be shown in supplementary material Table S1. We also used the dataset constructed by Kuang et al. [35] to further evaluate the pros and cons of the DNN-DTIs. The number of targets and drugs were 809 and 786, respectively, and the interacting DTIs are 3681. Finally, we utilize two DTIs network called Network1 and Network2 to to verify the effectiveness of DNN-DTIs. The Network1 was from Shi et al. [26], which was collected from [5], [6], and ChEMBL [36]. And the Network2 was from Xia et al. [24]. The numbers of the interaction pairs of the two networks are 33 and 150, respectively.

### 2.2. Protein feature extraction methods

#### 2.2.1. Composition, transformation and distribution

Composition (C), transition (T) and distribution (D) global protein descriptors are proposed by Dubchak et al. [37] in 1995, and CTD could describe and clarify the positions and physicochemical information of global and local amino acids. C represents the composition-based grouped amino acid information, T indicates the dipeptide-based frequency information, and D represents frequency and position information in the first, 25%, 50%, 75%, and the last occurrence of the grouped protein sequence. In this paper, thirteen physicochemical properties are used to group amino acids. Taking hydrophobicity as an example, the amino acid character signals are converted into three groups (PO (polar), NE (neutral) and HY (hydropphobic)). The calculation formulas of C, T, and D are shown in equation (1), (2) and (3):

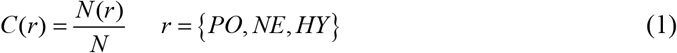

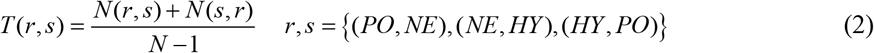

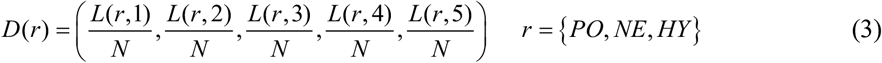

Where *N* (*r*) is the size of the *r*-*th* group in the amino acid. *N* is the sequence length. *N* (*r*, *s*) is the occurrence frequency of the dipeptide from *r*-*th* group to *s*-*th* group. *L*(*r*,1), *L*(*r*, 2), …, *L*(*r*,5) indicate the position information of the *r*-*th* group of amino acid at the first, 25%, 50%, 75% and 100%.

#### 2.2.2. Conjoint Triad

The conjoint triad (CT) was proposed by Shen et al. [38] in 2007. This method considers not only the composition pattern but also sequence order konwledge. First, 20 amino acids are grouped into seven categories. Then, the adjacent three residues are regarded as a whole block, and the frequency of conjoint triad can be calculated using equation (4):

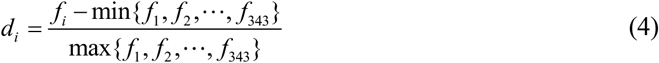

#### 2.2.3. Pseudo amino acid composition

Pseudo amino acid composition (PseAAC) [39] is a feature encoding method combining amino acid composition, physicochemical properties and statistical information (equation (5) and (6)).

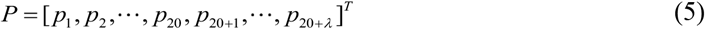

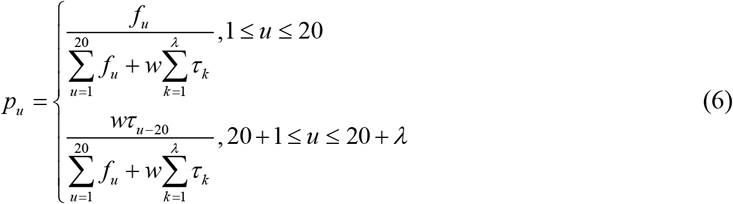

Where *p*_1_, *p*_2_, …, *p*_20_ indacates the amino acid composition information, and *p*_20+1_, … *p*_20+*λ*_ represents the sequence order information. *λ*(*λ* < *N*) reflects the sequence-related factors at different levels. *w* represents the weight factor.

#### 2.2.4. Pseudo position-specific scoring matrix

Pseudo position-specific scoring matrix (PsePSSM) is obtained through the PSSM matrix [40]. The PSI-BLAST program [41] is used in the Swiss-Prot database for comparison. After normalization, the PSSM is converted into PsePSSM feature vectors with the same length and biological significance through equations (8) and (9):

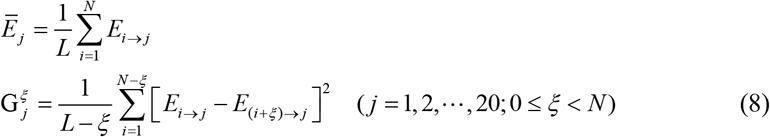

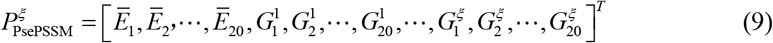

where 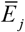 represents the average score at *j*-*th* position. 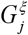 can illustrate the order evoluational information. *ξ* represents the interval.

#### 2.2.5. Normalized Moreau-Broto Autocorrelation

The Normalized Moreau-Broto (NMBroto) is a type of topology autocorrelation descriptor (equation (10) and (11)) proposed by Horne [42]. It is defined using the distribution of amino acid physicochemical properties.

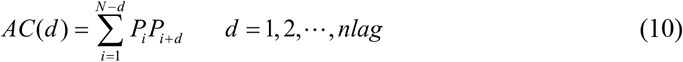

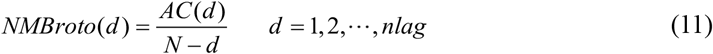

where, *d* is called the lag of autocorrelation (already known as the interval of amino acids), *P_i_* and *P_i_* + *d* are the amino acid property value at the position *i* and position *i* + *d*, *nlag*(*nlag* < *N*) is the maximum value of lag. This paper uses 8 types of physicochemical properties in the AAIndex database [43].

#### 2.2.6. Structural feature

This paper uses SPIDER3 [44] to generate SPD file (http://sparks-lab.org), which includes accessible surface area (ASA), secondary structure (SS), torsion angle (TA) and structural probability (SP) and etc. ASA is a *N* × 1 dimensional vector containing the accessible surface area values of all amino acid residues, and SS is also a *N* × 1 dimensional vector, including three types: *α*-helix (H), *β*-fold (E) and random coil (C). TA is a matrix with the size of *N* × 8, including the sine and cosine values of *φ*, *ψ*, *τ*, *θ*. SP is a matrix with size of *N* × 3 size. The obtained feature vectors can be analyzed via equations (12)–(17) proposed by Rayhan et al. [45].

1. Secondary Structure Composition

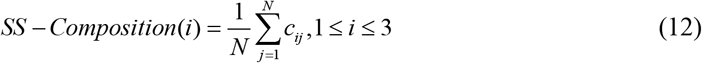

where *N* is the length of the protein sequence, 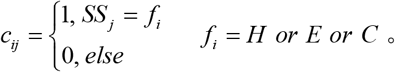
2. Accessible surface area composition

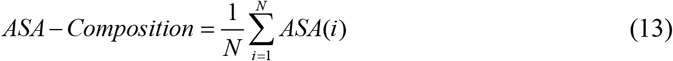
3. Torsional angles composition

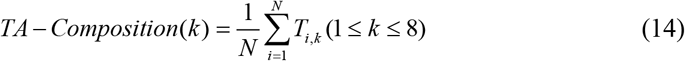
4. Torsional angles bigram

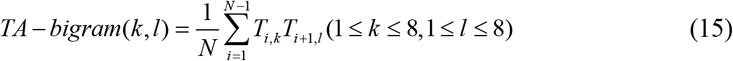
5. Structural probabilities bigram

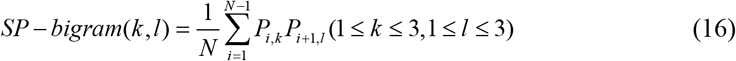
6. Torsional angles auto-covariance

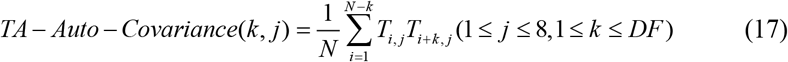

where *DF* is the distance factor whose value is 10 [46].
7. Structural Probablities Auto-Covariance

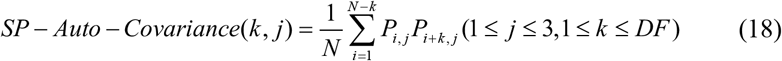

To sum up, for each feature extraction of target, the dimension of the obtained vectors are 273 (CTD), 343 (CT), (structure feature). 20+*λ* (PseAAC), 20 + 20 × *ξ* (PsePSSM), 8 × *nlag* (NMBroto), 195 (structure feature).

### 2.3. Drug molecular substructure fingerprint

Molecular substructure fingerprints can well characterize drug compounds, especially, the binary substructure vectors can well represent drug compound. By detecting the presence of specific molecular structure fragments in the molecular structure, drug can be converted into binary feature vector. This paper utilizes PaDEL-Descriptor [47] to calculate molecular fingerprint descriptors, describing the structure information by 881 chemical substructures collected from PubChem database. Therefore, if the substructure exists in drug molecule, the vectors of the corresponding position is set as 1. If the substructure does not exist, the value of this position is set as 0. These 881 chemical structure descriptions can be downloaded from (https://pubchem.ncbi.nlm.nih.gov), and the fingerprint attribute is “PUBCHEM_CACTVS_SUBGRAPHKEYS”.

### 2.4. XGBoost feature selection

XGBoost feature selection [48] is employed to calculate the feature importance score through gain (Equation (19)) to perform dimensionality reduction.

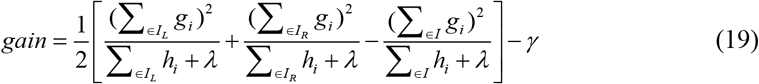

Where *I* = *I_L_* ⋃ *I_R_*, and *I_L_* is the number of sample in the left node space. *λ* and *γ* are regularization parameters. The higher the feature gain is, the more effective and important the feature is. The researcher can obtain XGBoost software from URL https://github.com/dmlc/xgboost.

### 2.5. Synthetic minority oversampling technique

Synthetic minority oversampling technique (SMOTE) [49], synthesizing new samples to balance the samples, which could deal with the imbalanced prediction problem (The number of DTIs samples is larger than that of non-DTIs samples). The SMOTE can be obtained using equation (20). First, the *k* nearest neighbors are determined by calculating the Euclidean distance. And the random netighbor *x_n_* can be determined through the sample imbalance ratio *N*. Then the new samples can be obtained combining the original sample according to equation (20).

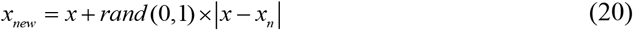

### 2.6. Deep neural network

Deep neural network (DNN) was presented by Hinton et al. [50] using the idea of human brain learning. The input layer receives raw sample vectors. Each value multiplies by the weights at each node, and the output values can be obtained though the nonlinear activation function. The weights and biases can be adjusted using stochastic gradient descent (Fig. 1 (A)). The mathematical representations are shown in equation (21).

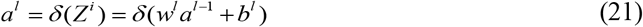

Where *l* = 1, 2, ···, *N*, *a^l^* represents the input data of the layer, *w^l^* is the connection weight matrix, *b^l^* is the bias of the layer, and *δ* represents the activation function of the *l*-*th* layer. DNN-DTIs uses the cross-entropy loss function for optimization:

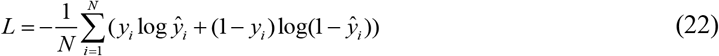

where *N* is sample number, *y_i_* and 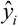 represent the true label and predictive label, respectively.

**Fig. 1.**
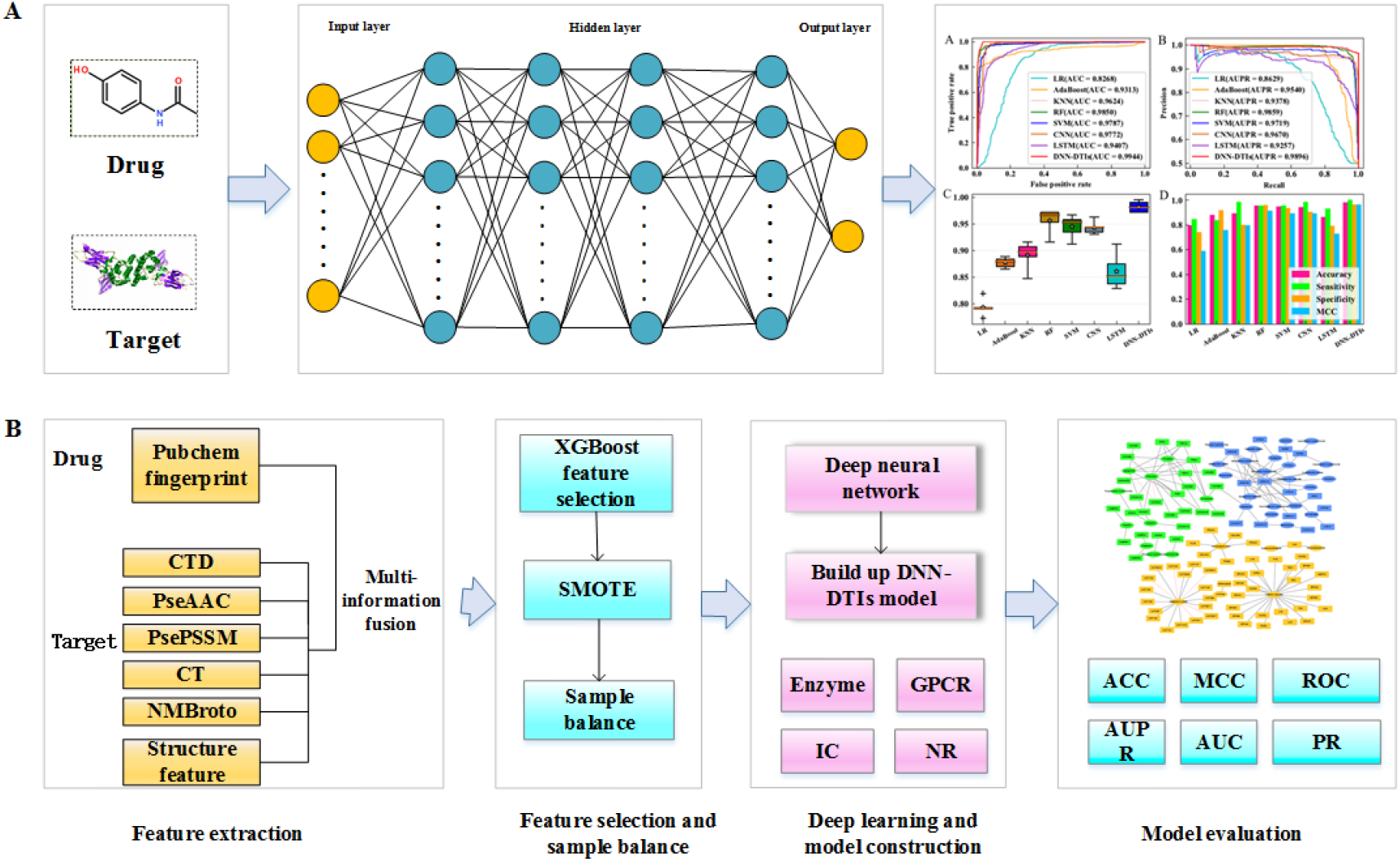
DTIs prediction method based on deep neural network. (A) is the structure of the deep neural network and its inputs and outputs. The input should be drug and protein, and the output can be prediciton labels and evaluation indicators. (B) is the pipeline of DNN-DTIs. This paper fuses CTD, PseAAC, PsePSSM, CT, NMBroto and structure feature to represent target information. The drug information is represented using fingerprint. Then XGBoot and SMOTE are used to process the tasks of feature selection and sample balance. On Enzyme, GPCR, IC and NR, the DNN-DTIs model is built up. Finally, the DTIs networks are predicted and ploted.

ReLU (Rectified linear unit) has one-sided suppression characteristics, thus, all negative values are changed to 0, while positive values are unchanged (equation (23)).

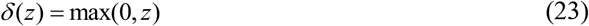

For this DL framework, the input layer consists of extracted features, and the hidden layer has 4 layers (the numbers of neurons are 600, 300, 150, and 75, respectively). The activation function is set to ReLU, and the two neurons in the output layer represent the output probabilities of DTI and non-DTI, respectively. We use Adam algorithm to optimize the model.

### 2.7. Performance evaluation and model construction

In order to evaluate the effectiveness of the model, this paper uses 5-fold cross-validation (5-fold CV) for evaluation. Sensitivity (SE), Specificity (SP), Matthew’s correlation coefficient (MCC) and accuracy (ACC) are used to evaluate DNN-DTIs [51,52], which can be find in equation (24), (25), (26) and (27).

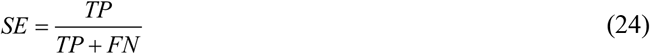

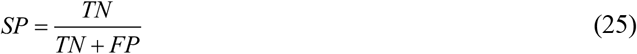

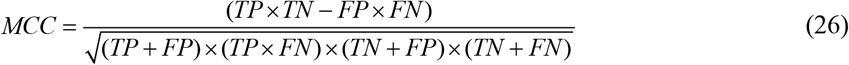

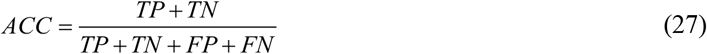

where *TP*, *FP*, *TN* and *FN* are true positive, false positive, true negative and false negative, respectively. In addition, ROC and PR curve are also important indicators and the areas under the curves are called AUC and AUPR, respectively. The larger the value is, the better the performance of DNN-DTIs is.

The novel prediction method of DTIs is called DNN-DTIs, whose calculation pipeline is shown in Fig. 1. The experimental experiments are conducted on Intel Xeon (E5-2650) 32.0GB of memory and Python programming. The code and dataset of DNN-DTIs are available on https://github.com/QUST-AIBBDRC/DNN-DTIs.

The calculational process of DNN-DTIs can be described as:

1. Feature extraction. Physicochemical information, local and global information of sequence composition, and structural information of protein sequences are obtained through CTD, PseAAC, PsePSSM, CT, Moreau-Broto and structure feature. The chemical structure information of the drug can be characterized using molecular substructure fingerprint.
2. Feature selection and sample balance. For eliminating noise information, the importance score of each feature is calculated by XGBoost, and then the low-dimensional and useful essential features are obtained. The XGBoost is compared with information gain, Gini index, max-relevance-max-distance, LASSO and elastic net. The SMOTE is employed to balance the samples of DTIs and non-DTIs, generating a balanced and effective DTIs dataset and boosting generalization performance.
3. Deep learning and model construction. The drug-target interactions prediction model DNN-DTIs based on deep neural network is constructed through determining various parameters. Compared with CNN, long short-term memory neural network (LSTM) and traditional machine learning methods, DNN-DTIs obtains better performance on Enzyme, IC, GPCR, NR, and Kuang’s dataset.
4. In order to further evaluate the pros and cons of DNN-DTIs, the validation is verified on Network1 and Network2. Then the DTIs networks are ploted to analyze the network topology and explore the biomedical significance.

## 3. Results and discussion

### 3.1. Parameter optimization of DNN-DTIs model

Feature extraction, converting protein character signals into numerical signals, can mine physicochemical, local and global sequence, structural and evolutional information of the target. In this paper, the parameter *λ* of PseAAC, *ξ* of PsePSSM, and *nlag* of Moreau-Broto autocorrelation could generate influence to the DNN-DTIs model performance. Considering the shortest sequence length is 83 in gold standard datasets, this paper sets different values to optimize the parameters based on DNN (one layer of input layer, four hidden layers and one output layer). The results on GSD via 5-fold CV are shown in Table S3-S5.

For PseAAC, the highest accuracy are reached when *λ* = 30, *λ* = 70, *λ* = 70 and *λ* = 80 on Enzyme, IC, GPCR and NR, respectively (Table S3). Table S4 shows that the highest accuracy is achieved when using *ξ* = 0, *ξ* = 6, *ξ* = 6 and *ξ* = 8 on Enzyme, IC, GPCR and NR, respectively. From Table S5, the average prediction accuracy of GSD is the greatest when *nlag* = 6. According to Table S3-S5, the performance of different single feature change, especially, the accuracies of the enzyme vary greatly, while the results of IC, GPCR and NR changes little. Finally, the optimal parameters of PseAAC, PsePSSM and Moreau-Broto autocorrelation are *λ* = 30, *ξ* = 2 and *nlag* = 6, respectively.

### 3.2. Comparison results of feature fusion with single feature

Extracting effective information from protein sequences is critical for DTIs prediction. In this paper, the CTD and CT are adopted from the perspective of amino acid sequence composition, and then PseAAC and Moreau-Broto are employed to mine physicochemical information. PsePSSM is used to extract evolutionary information. Finally we fuse multiple information to characterize comprehensive and effective DTIs feature information. According to section 3.1, the number of dimension after feature fusion is 1850. The comparasion results of feature fusion with single feature through 5-fold cross-validation are shown in Table 1. The ROC and PR curves of the individual feature and multi-feature fusion are shown in Fig. S1. The vectors of each feature encoding algorithm is processed using SMOTE. The DNN is utilized as a classifier, and Fusion represents the result of multi-feature fusion.

**Table 1.**
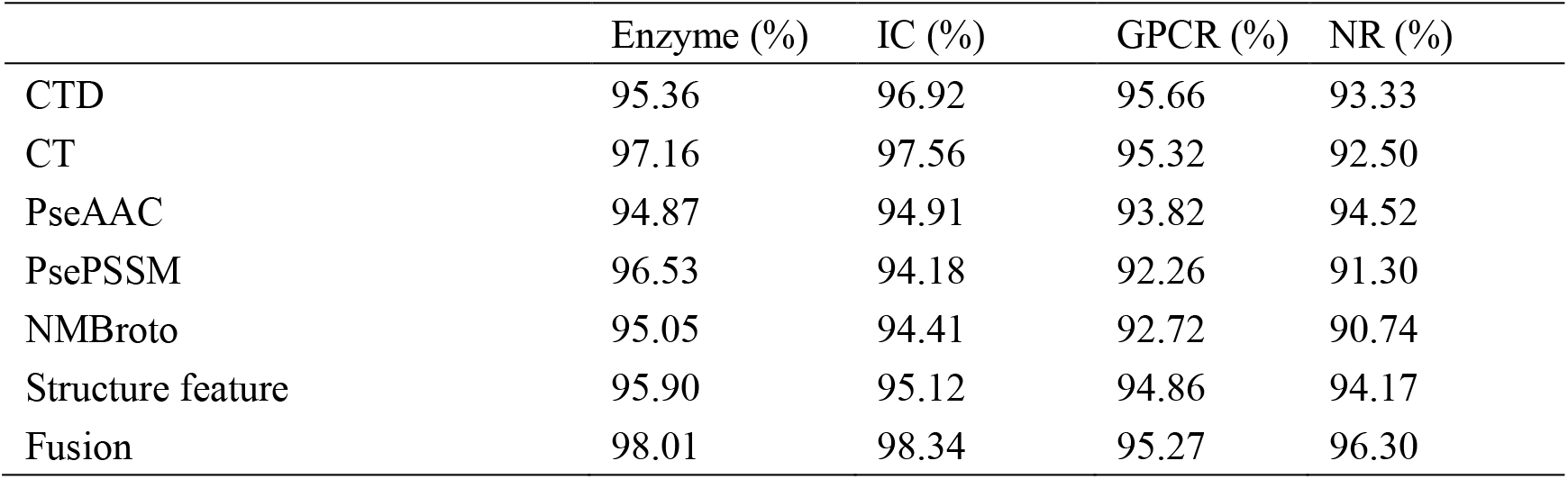
Comparison of single feature and feature fusion.

|  | Enzyme (%) | IC (%) | GPCR (%) | NR (%) |
| --- | --- | --- | --- | --- |
| CTD | 95.36 | 96.92 | 95.66 | 93.33 |
| CT | 97.16 | 97.56 | 95.32 | 92.50 |
| PseAAC | 94.87 | 94.91 | 93.82 | 94.52 |
| PsePSSM | 96.53 | 94.18 | 92.26 | 91.30 |
| NMBroto | 95.05 | 94.41 | 92.72 | 90.74 |
| Structure feature | 95.90 | 95.12 | 94.86 | 94.17 |
| Fusion | 98.01 | 98.34 | 95.27 | 96.30 |

Table 1 shows that different special extraction methods make the prediction accuracy of DTIs different. The result after fusion of multiple information is better than the prediction result of single feature. In addition to the GPCR dataset, ACC of CTD exceeds Fusion. For the other three datasets, Fusion’s prediction performance is the best, indicating that feature fusion can effectively characterize drugs and targets. Figure S1 indicates that the ROC using the Fusion has the largest coverage area, whose AUC values are 0.9969, 0.9965, 0.9836, and 0.9858 on four GSD, respectively. Therefore, the prediction performance using multiple information fusion is the best and has good robustness.

### 3.3. XGBoost feature selection

When the XGBoost algorithm for feature selection is used, different dimensions have an important impact on the prediction results. Fusing CTD, CT, PseAAC, PsePSSM, NMBroto and structure feature can construct raw feature space with *λ*=30 、 *ξ* = 2 and *nlag* = 6. We utilize XGBoost and SMOTE algorithm to select feature and balance dataset, and deep neural network is employed to predict DTIs. In order to determine the optimal and most effective number of feature subsets, we set the size of the feature subsets to 100, 200, 300, 400, 500, 600, 700, 800, 900, and 1000 in turn. The main results of DNN-DTIs can be find form Table S6-Table S9, and ROC and PR curves are shown in Fig. 3 with different dimensions. Fig. 3 and Table S6-Table S9 show that XGBoost feature selection is employed to perform dimension reduction with the dimension of 300, making the performance of DNN-DTIs is optimal. The optimiazed features can provide high-level feature information representing DTIs for deep learning. When the dimension is determined to be 1000, the prediction results of the four gold standard datasets are the worst. With the number of subsets increases, the prediction accuracy rate gradually decreases. So the 300 optimal features could better represent DTIs and offer effective information for DNN.

XGBoost, a tree-based feature selection method, can mine the nonlinear relationship between DTIs features and label and preserve effective feature subset. We tested the impact of other feature selection methods on the prediction of DTIs. For information gain [53] (IG), the number of retained best feature subsets is set to 300. For GINI Index [54] (GINI), the number of retained optimal feature subsets is set to 300. For max-relevance-max-distance [55] (MRMD), the optimal subset number is set to 300. For LASSO [56], The penalty parameter alpha is set to 0.01. And for the elastic net [57] (EN), the penalty parameters alpha and l1_ratio are set to 0.1 and 0.05. The prediction results and evaluation on Enzyme, IC, GPCR and NR are shown in Table 2 and Fig. 2. The optimal dimensions of different feature selection can be find fromTable S10. We also obtain the comparison results of different feature selection methods in the dataset constructed by Kuang et al. [35] (Table S11), the ROC curve and the PR curve are shown in Fig. S3.

**Fig. 2.**
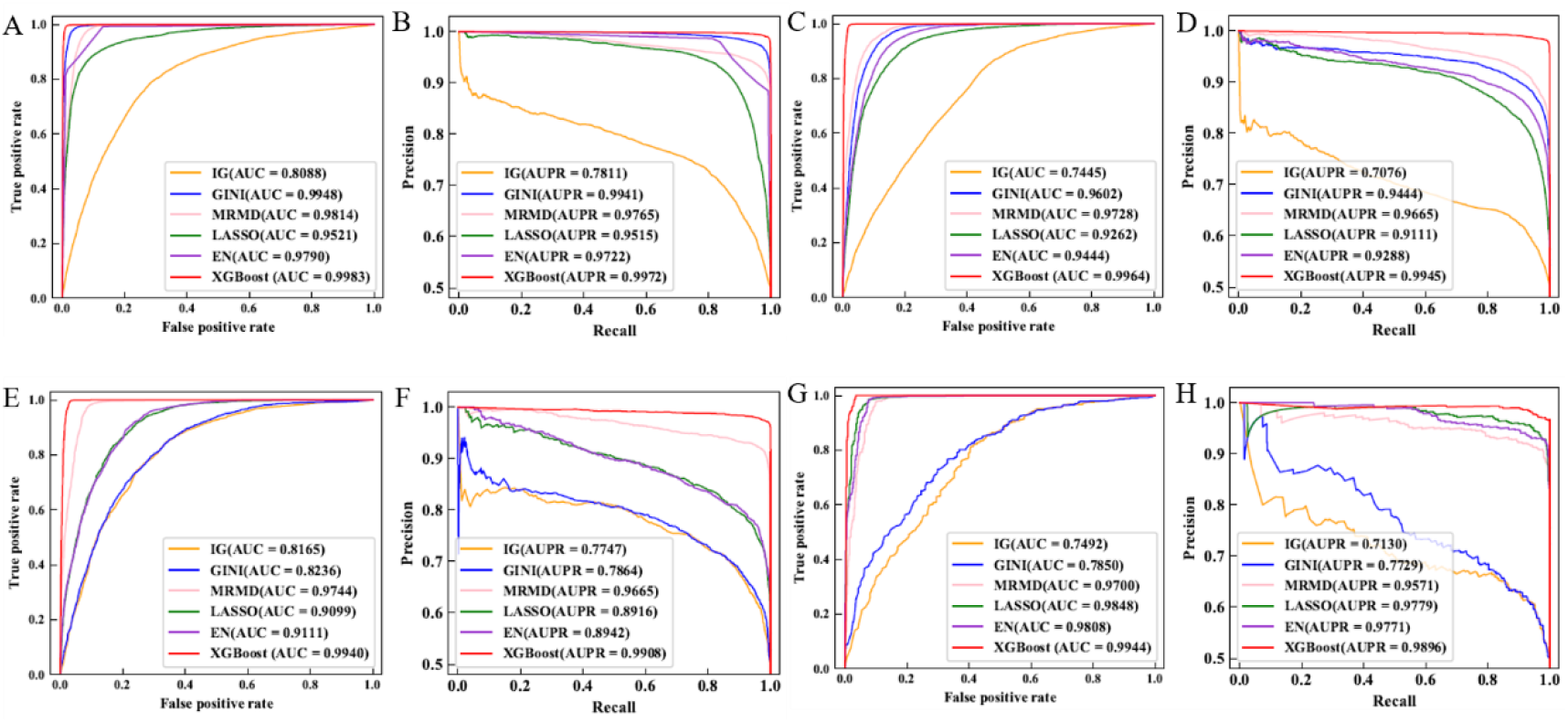
ROC and PR curves of the gold standard datasets under IG, GINI, MRMD, LASSO, EN and XGBoost feature selection methods. (A-B), (C-D), (E-F) and (G-H) are ROC and PR curves of Enzyme, IC, GPCR and NR, respectively.

**Table 2.**
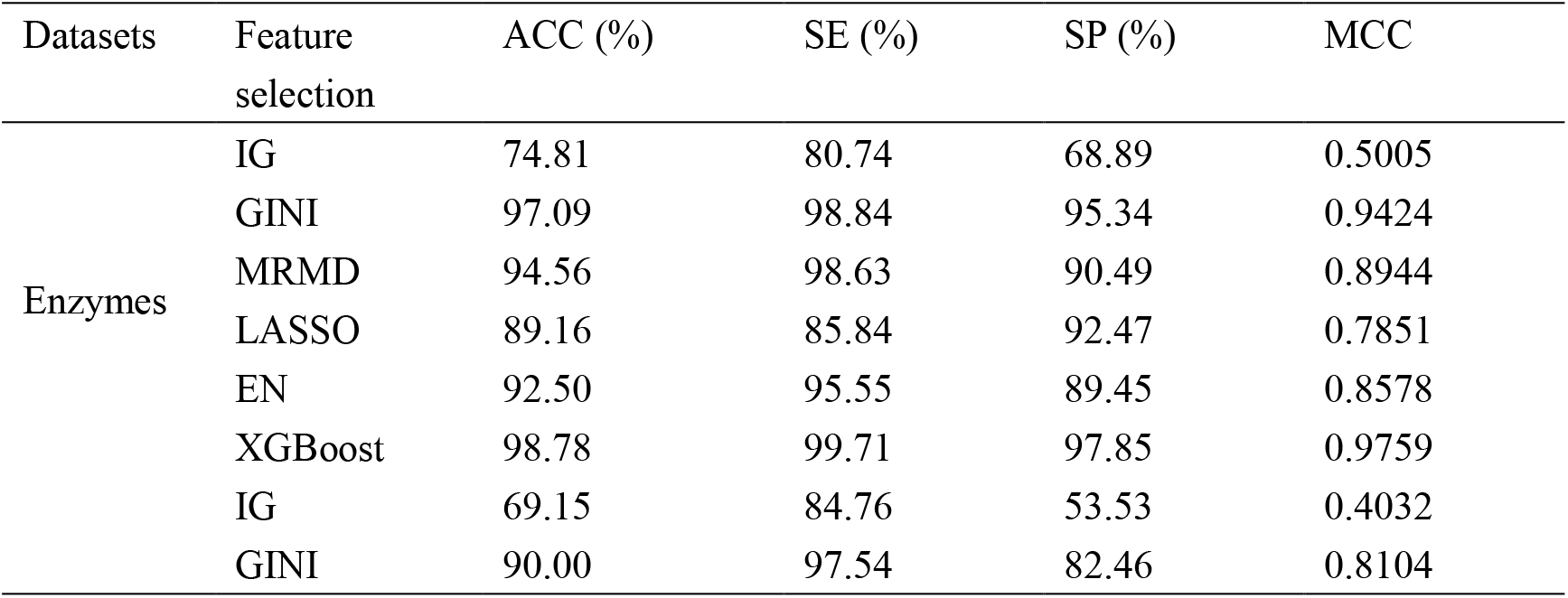
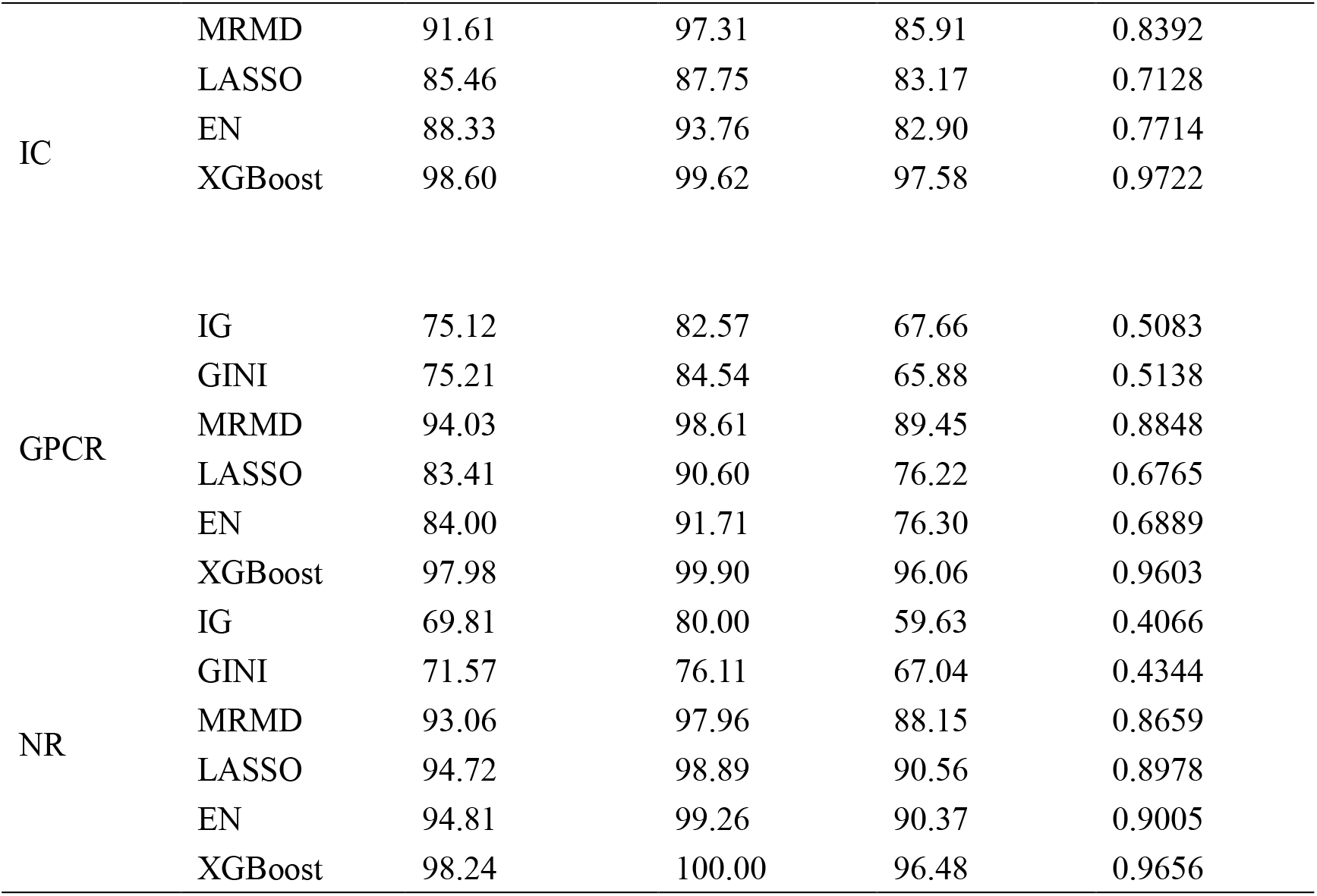
Comparison results of XGBoost and other feature selection methods on the gold standard dataset.

As shown in Table 2, XGBoost feature selection methods all exceed the model prediction performance of information gain (IG), Gini coefficient (GINI), maximum correlation maximum distance (MRMD), LASSO and elastic net (EN). For Enzyme, XGBoost is 23.97% higher than IG (98.78% VS 74.81%) on ACC and 9.62% higher than LASSO (98.78% VS 89.16%). For the IC dataset, XGBoost’s ACC values are 29.45%, 8.6%, 6.99%, 13.14%, and 10.27% higher than IG, GINI, MRMD, LASSO, and EN (98.6% VS 69.15%, 90%, 91.61%, 85.46%, 88.33%), and MCC value is 13.3% higher than MRMD (0.9722 VS 0.8392). For GPCR, the ACC value of XGBoost is 22.86%, 22.77%, 3.95%, 14.57%, 13.98% higher than IG, GINI, MRMD, LASSO and EN (97.98% VS 75.12%, 75.21%, 94.03%, 83.41% and 84%.). In terms of MCC value, XGBoost is 45.2% higher than IG (0.9603 VS 0.5083). For the NR, XGBoost’s ACC values are 28.43%, 26.67%, 5.18%, 3.52%, and 3.43% higher than IG, GINI, MRMD, LASSO, and EN (98.24% VS 69.81%, 71.57%, 93.06%, 94.72%, 94.81%). For MCC, the XGBoost outperform LASSO (0.9656 VS 0.8978) and EN (0.9656 VS 0.9005).

Figure 2 (A) shows that for Enzyme, the AUC of XGBoost is 18.95%, 0.35%, 1.69%, 4.62%, 1.93% (0.9983 VS 0.8088, 0.9948, 0.9814, 0.9521, 0.979) higher than IG, GINI, MRMD, LASSO and EN). On AUPR, XGBoost is 21.61% better than IG (0.9972 VS 0.7811) and 4.57% better than LASSO (0.9972 VS 0.9515). It can be seen from Figure 2 (B). For IC, XGBoost’s AUC values are 25.19%, 3.62%, 2.36%, 7.02%, and 5.2% higher than IG, GINI, MRMD, LASSO, and EN (0.9964 VS 0.7445, 0.9602, 0.9728, 0.9262, 0.9444). Figure 2 (C) indicates that on the GPCR dataset, the AUC values of XGBoost is 17.75%, 17.04%, 1.96%, 8.41%, 8.29% (0.994 VS 0.8165, 0.8236, 0.9744, 0.9099, 0.9111). Figure 2 (D) indicates that for NR, the AUC value of XGBoost is 24.52%, 20.94%, 2.44%, 0.96%, 1.36% higher than IG, GINI, MRMD, LASSO, and EN (0.9944 VS 0.7492, 0.785, 0.97, 0.9848, 0.9808). XGBoost’s AUPR values are 27.66%, 21.67%, 3.25%, 1.17%, 1.25% higher than other feature selection approaches (0.9896 VS 0.713, 0.7729, 0.9571, 0.9779, 0.9771).

From the above analysis, XGBoost feature selection methods all exceed the predicted performance of IG, GINI, MRMD, LASSO, and EN. XGBoost can calculate the variable score based on the tree structure and boosting algorithm to determine the important feature subset with a large contribution rate, which removes the noise information and redundancy that is not related to DTIs prediction. In other words, XGBoost can reduce the computational complexity and improve the operating efficiency. Other feature selection methods have their pros and cons, and XGBoost is superior to information theory-based feature selection methods (IG, GINI, and MRMD) and regularization-based feature selection methods (LASSO and EN) for prediction of DTIs. Therefore, XGBoost feature selection method is used in this paper to perform dimensionality reduction.

### 3.4. Comparison results of DNN with other classifiers

For studying the contribution of deep neural networks to predicting DTIs and its powerful hierarchical learning ability, we complete feature extraction, feature selection and imbalance processing. The low-dimensional and effective feature information is input into the classifiers, including logistic regression [58] (LR), AdaBoost [59] (the number of base decision trees is 500), K nearest neighbor algorithm [60] (KNN, neighbor size is set to 5), random forest [61] (RF, the value of n_estimators is set to 500), support vector machine [62] (SVM, using radial basic kernel function), convolutional neural network [63] (CNN, including convolutional layer, pooling layer, activation function is rectification function), long and short term memory neural network [64] (LSTM, the activation function is a rectification function). The main prediction results of GSD can be obained from Table 3-Table 6. In order to further intuitively express comparison results with other ML and DL methods, ROC curve, PR curve, box chart and histogram are shown in Fig. 3-Fig. 6.

**Fig. 3.**
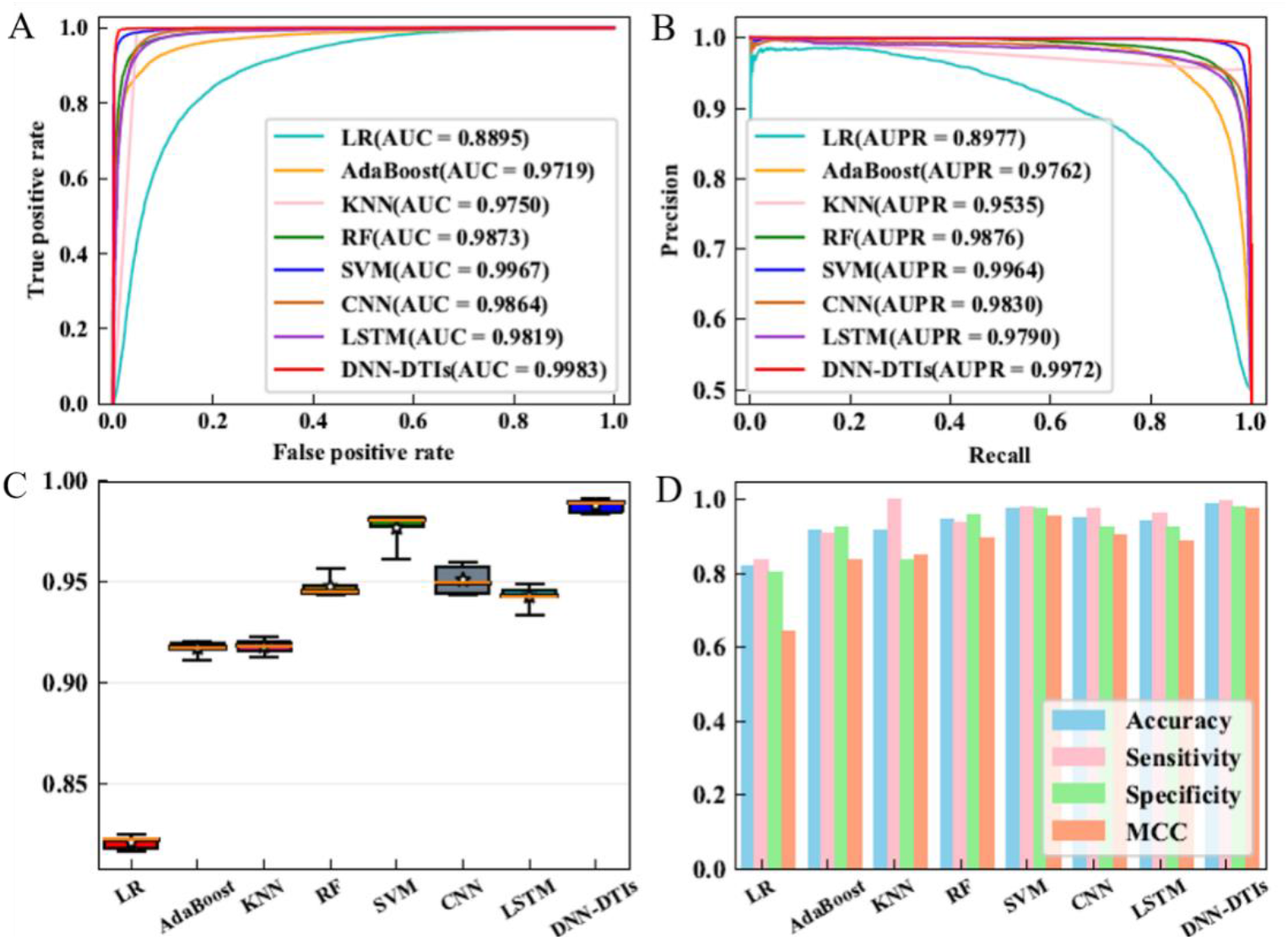
ROC curves (A), PR curves (B), box plots (C) and histograms (D) of LR, AdaBoost, KNN, RF, SVM, CNN, LSTM and DNN-DTIs on Enzyme dataset.

**Table 3.** The main results of different classifiers for Enzyme dataset.

| Methods | ACC (%) | SE (%) | SP (%) | MCC | AUC |
| --- | --- | --- | --- | --- | --- |
| LR | 82.08 | 83.75 | 80.40 | 0.6420 | 0.8895 |
| AdaBoost | 89.51 | 86.16 | 92.87 | 0.8077 | 0.9719 |
| KNN | 91.79 | 99.84 | 83.74 | 0.8469 | 0.9750 |
| RF | 94.59 | 93.29 | 95.88 | 0.8929 | 0.9873 |
| SVM | 97.68 | 97.92 | 97.43 | 0.9538 | 0.9967 |
| CNN | 95.10 | 97.63 | 92.56 | 0.9033 | 0.9864 |
| LSTM | 94.28 | 96.24 | 92.32 | 0.8864 | 0.9819 |
| DNN-DTIs | 98.78 | 99.71 | 97.85 | 0.9759 | 0.9983 |

From Table 3 to Table 6, we can see that the prediction results of DNN-DTIs are better than the other 7 classifiers. Table 4 shows that for the Enzyme dataset, the ACC values of DNN-DTIs are 16.7%, 9.27%, 6.99%, 4.19%, 1.1%, 3.68% and 4.5% higher than LR, AdaBoost, KNN, RF, SVM, CNN, and LSTM, respectively (98.78% VS 82.08%, 89.51%, 91.79%, 94.59%, 97.68%, 95.10% and 94.28%). Table 5 shows that for the IC, the ACC values of DNN-DTIs are 14.98%, 10.41%, 8.44%, 5.81%, 1.6%, 4.36%, 4.98% and 9.05% higher than LR, AdaBoost, KNN, RF, SVM, CNN, and LSTM, respectively (98.60% VS 83.62%, 88.19%, 90.16%, 92.79%, 97.00%, 94.24% and 89.55%). Table 6 shows that for GPCR, the ACC values of DNN-DTIs are 18.08%, 9.71%, 7.95%, 4.62%, 1.64%, 4.25%, 3.8% and 12.46% higher than LR, AdaBoost, KNN, RF, SVM, CNN, and LSTM, respectively (97.98% VS 79.90%, 88.27%, 90.03%, 93.36%, 96.34%, 93.73%, 85.52%). Table 7 indicates that for NR, the ACC values of DNN-DTIs are 18.7%, 11.39%, 9.44%, 3.43%, 4.26%, 3.89% and LR, AdaBoost, KNN, RF, SVM, CNN and LSTM, respectively. 12.13% (98.24% VS 79.54%, 86.85%, 88.80%, 94.81%, 93.98%, 94.35% and 86.11%).

**Table 4.** The main results of different classifiers for IC dataset.

| Methods | ACC (%) | SE (%) | SP (%) | MCC | AUC |
| --- | --- | --- | --- | --- | --- |
| LR | 83.62 | 87.64 | 79.61 | 0.6747 | 0.8873 |
| AdaBoost | 88.19 | 84.84 | 91.53 | 0.7831 | 0.9661 |
| KNN | 90.16 | 99.30 | 81.02 | 0.8170 | 0.9657 |
| RF | 92.79 | 94.80 | 90.79 | 0.8592 | 0.9815 |
| SVM | 97.00 | 97.73 | 96.27 | 0.9404 | 0.9926 |
| CNN | 94.24 | 96.48 | 91.99 | 0.8861 | 0.9782 |
| LSTM | 89.55 | 94.77 | 84.33 | 0.7955 | 0.9546 |
| DNN-DTIs | 98.60 | 99.62 | 97.58 | 0.9722 | 0.9964 |

**Table 5.** The main results of different classifiers for GPCR dataset.

| Methods | ACC (%) | SE (%) | SP (%) | MCC | AUC |
| --- | --- | --- | --- | --- | --- |
| LR | 79.90 | 83.67 | 76.12 | 0.5999 | 0.8676 |
| AdaBoost | 88.27 | 83.83 | 92.70 | 0.7899 | 0.9591 |
| KNN | 90.03 | 99.27 | 80.79 | 0.8146 | 0.9556 |
| RF | 93.36 | 95.22 | 91.50 | 0.8707 | 0.9833 |
| SVM | 96.34 | 97.11 | 95.56 | 0.9274 | 0.9914 |
| CNN | 93.73 | 96.51 | 90.94 | 0.8763 | 0.9756 |
| LSTM | 85.52 | 91.08 | 79.97 | 0.7154 | 0.9273 |
| DNN-DTIs | 97.98 | 99.90 | 96.06 | 0.9603 | 0.9940 |

**Table 6.** The main results of different classifiers for NR dataset.

| Methods | ACC (%) | SE (%) | SP (%) | MCC | AUC |
| --- | --- | --- | --- | --- | --- |
| LR | 79.54 | 85.56 | 73.52 | 0.5972 | 0.8268 |
| AdaBoost | 86.85 | 84.07 | 89.63 | 0.7549 | 0.9313 |
| KNN | 88.80 | 98.33 | 79.26 | 0.7909 | 0.9624 |
| RF | 94.81 | 94.26 | 95.37 | 0.9010 | 0.9850 |
| SVM | 93.98 | 93.52 | 94.44 | 0.8810 | 0.9787 |
| CNN | 94.35 | 98.33 | 90.37 | 0.8906 | 0.9772 |
| LSTM | 86.11 | 92.96 | 79.26 | 0.7293 | 0.9407 |
| DNN-DTIs | 98.24 | 100.00 | 96.48 | 0.9656 | 0.9944 |

**Table 7.**
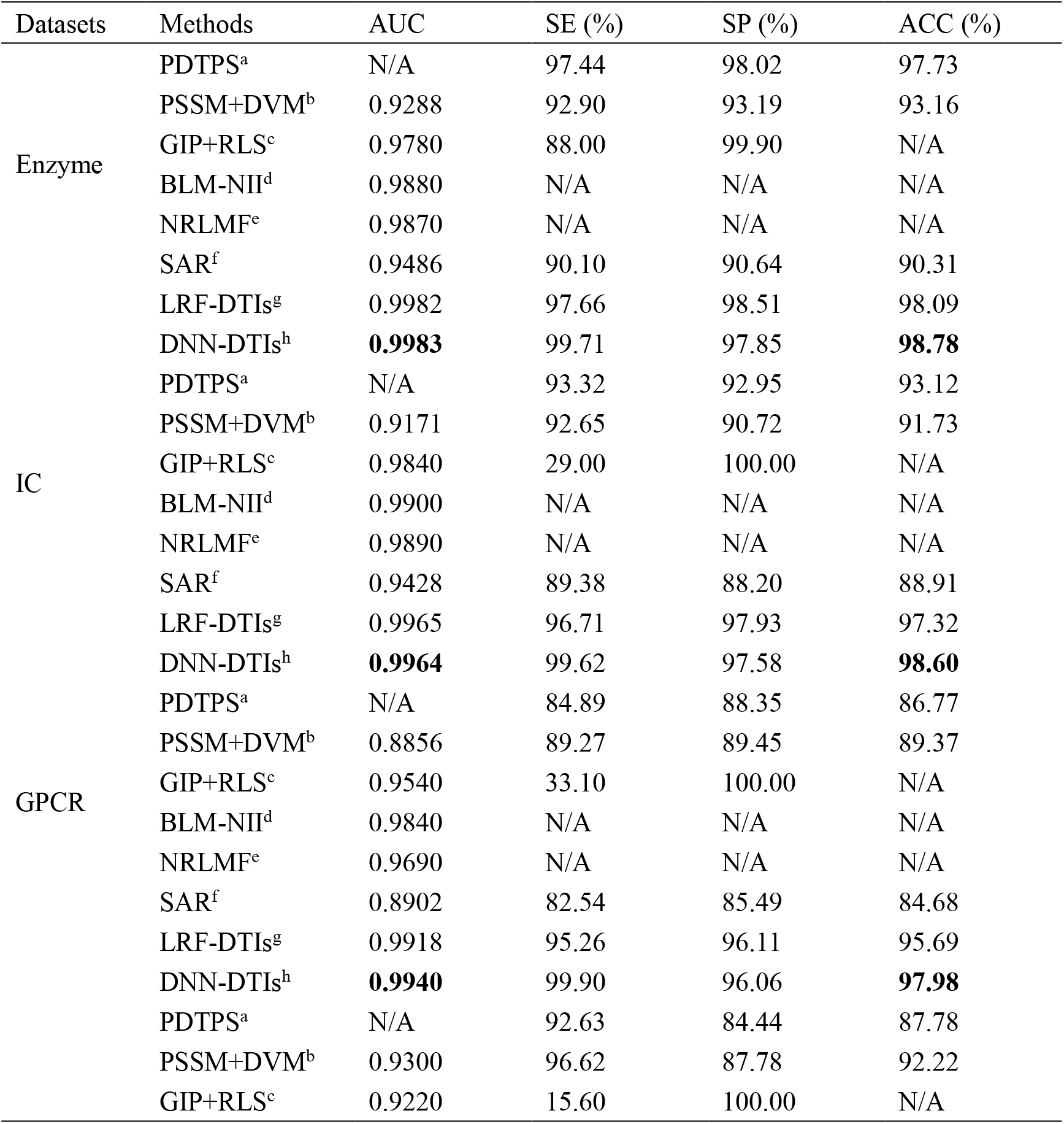
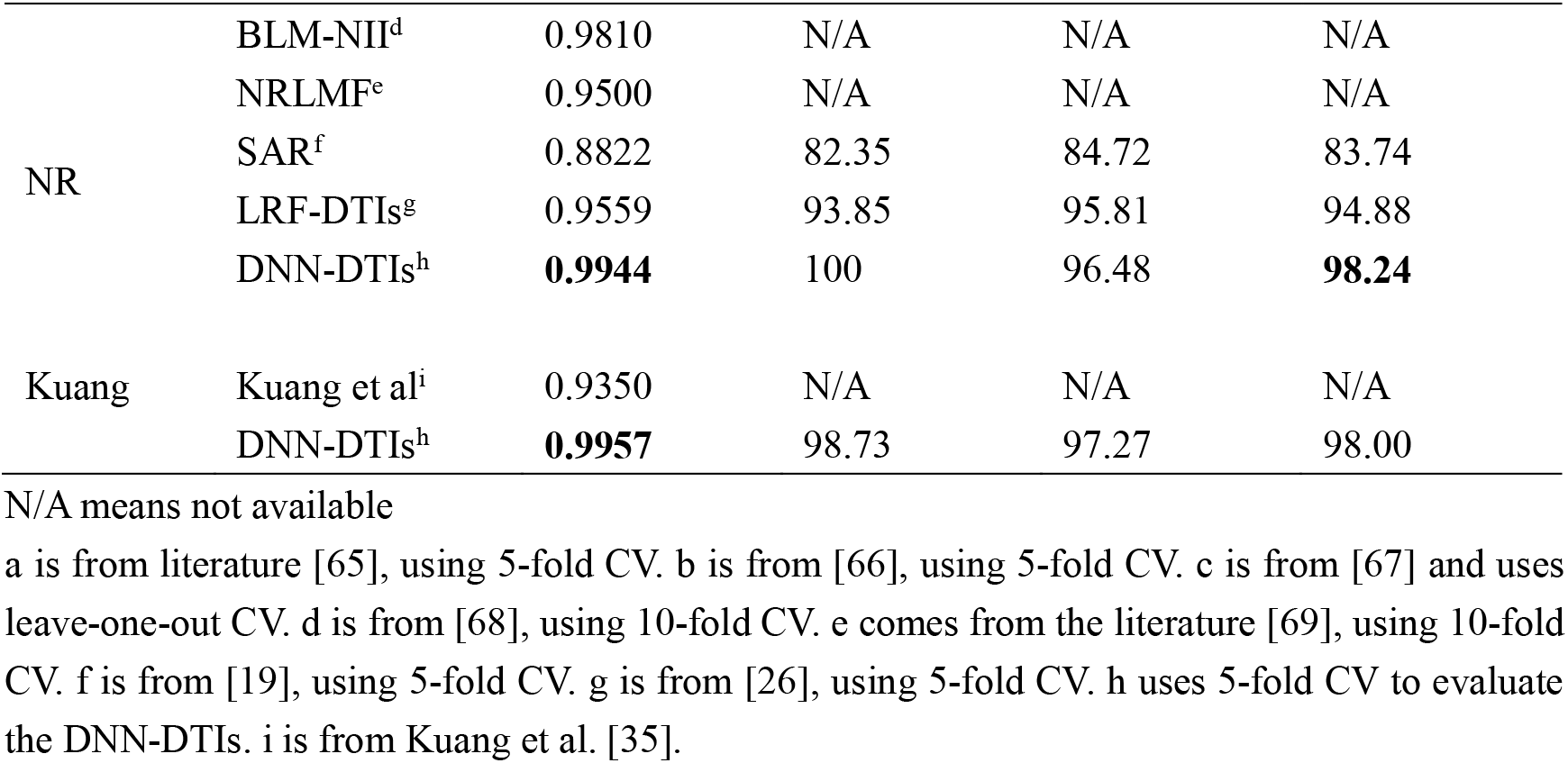
Comparison of different DTIs prediction methods on GSD and Kuang dataset.

Fig. 3 indicates that for Enzyme, from the ROC curve, PR curve, box chart and bar chart, DNN-DTIs ahieves better performance than that of LR, AdaBoost, KNN, RF, SVM, CNN and LSTM effect. The AUC value of DNN-DTIs is 0.9983, which is 1.19% better than CNN (0.9983 VS 0.9864) and 0.64% better than LSTM (0.9983 VS 0.9819). The AUC values of DNN-DTIs are 10.88%, 2.64%, 2.33%, and 0.16% higher than LR, AdaBoost, KNN, and SVM (0.9983 VS 0.8895, 0.9719, 0.975, and 0.9967). It can also be seen from the box plot and histogram that the robustness and prediction effect of DNN-DTIs are superior. Fig. 4 indicates that for IC, the AUC values of DNN-DTIs are 10.91%, 3.03%, 3.07%, 1.49%, 0.38%, 1.82%, 10.91% and 4.18% higher than LR, AdaBoost, KNN, RF, SVM, CNN, and LSTM, respectively (0.9964 VS 0.8873, 0.9661, 0.9657, 0.9815, 0.9926, 0.9782 and 0.9546). The AUPR value of DNN-DTIs is 1.42% higher than CNN (0.9972 VS 0.9830) and 0.96% higher than RF (0.9972 VS 0.9876). Fig. 5 shows that for the GPCR dataset, the AUC values of DNN-DTIs are 12.64%, 3.49%, 3.84%, 1.07%, 0.26%, 1.84%, 12.64% and 6.67% higher than LR, AdaBoost, KNN, RF, SVM, CNN, and LSTM, respectively (0.994 VS 0.8676, 0.9591, 0.9556, 0.9833, 0.9914, 0.9756 and 0.9273). The AUPR value of DNN-DTIs is also better than other classifier algorithms, and the box graph and histogram of GPCR can also illustrate its superiority intuitively. Fig. 6 shows that for NR, the AUC values of DNN-DTIs are 16.76%, 6.31%, 3.2%, 0.94%, 1.57%, 1.72%, 7.6% and 5.37% higher than LR, AdaBoost, KNN, RF, SVM, CNN, and LSTM, respectively. (0.9944 VS 0.8268, 0.9313, 0.9624, 0.985, 0.9787, 0.9772 and 0.9407). From Figure 9 (B) we can see that the area of the PR curve of DNN-DTIs is higher than other classifiers, which is 2.26% higher than CNN (0.9896 VS 0.9670). Figure 6 (C) and (D) also intuitively show that the prediction effect is better than other classifiers in terms of the main indicators.

**Fig. 4.**
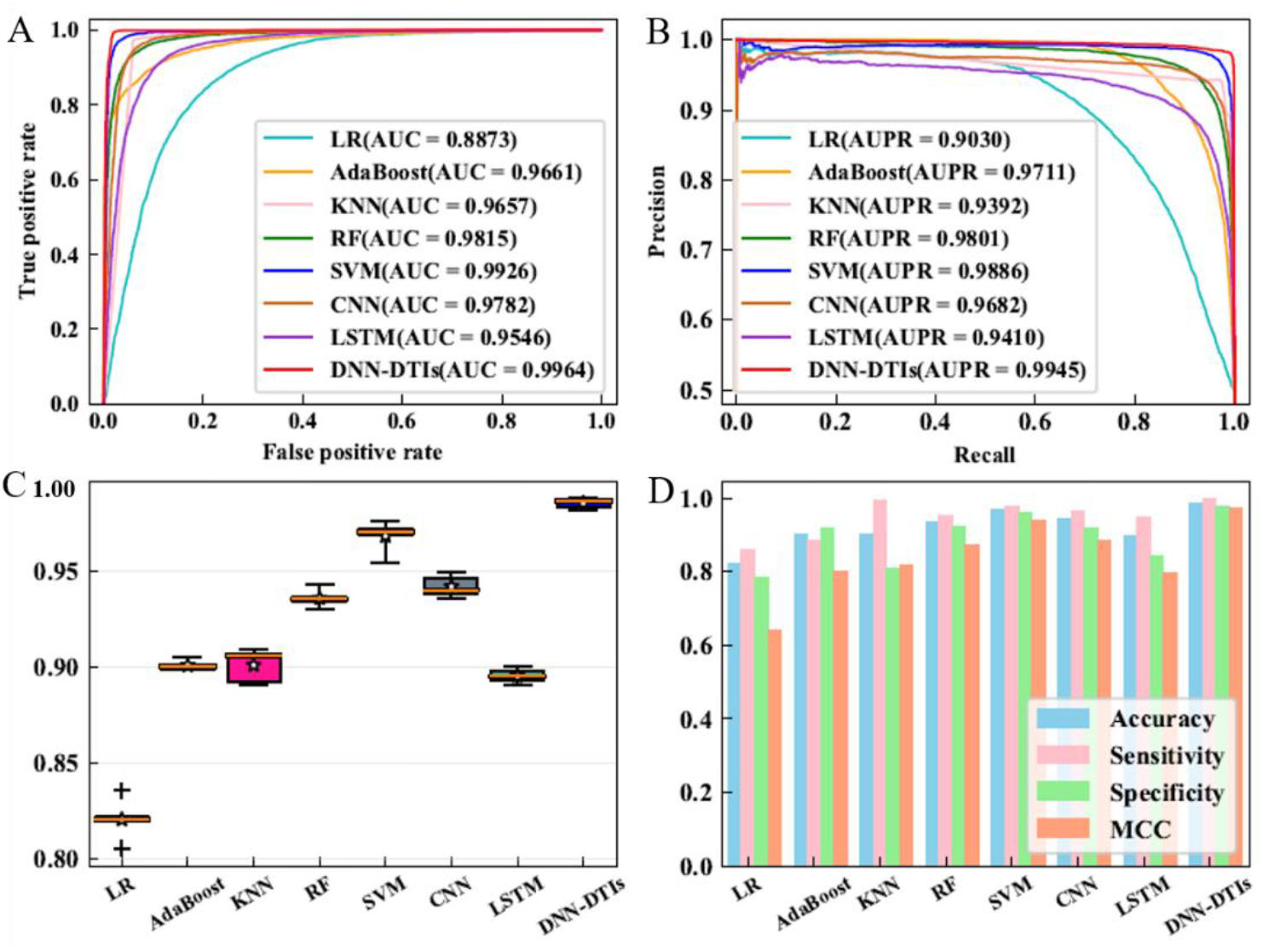
ROC curves (A), PR curves (B), box plots (C) and histograms (D) of LR, AdaBoost, KNN, RF, SVM, CNN, LSTM and DNN-DTIs on IC dataset.

**Fig. 5.**
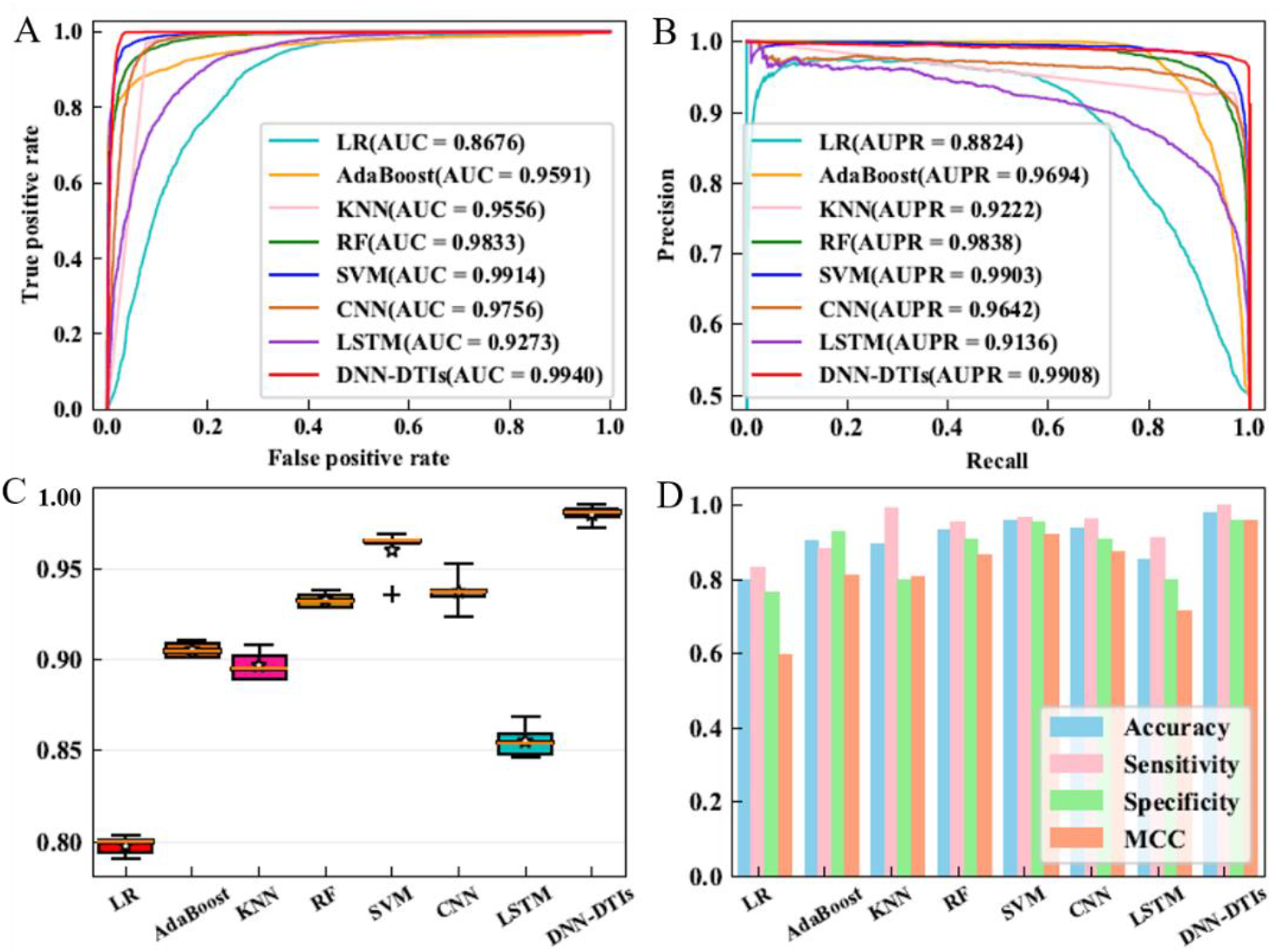
ROC curves (A), PR curves (B), box plots (C) and histograms (D) of LR, AdaBoost, KNN, RF, SVM, CNN, LSTM and DNN-DTIs on GPCR dataset.

**Fig. 6.**
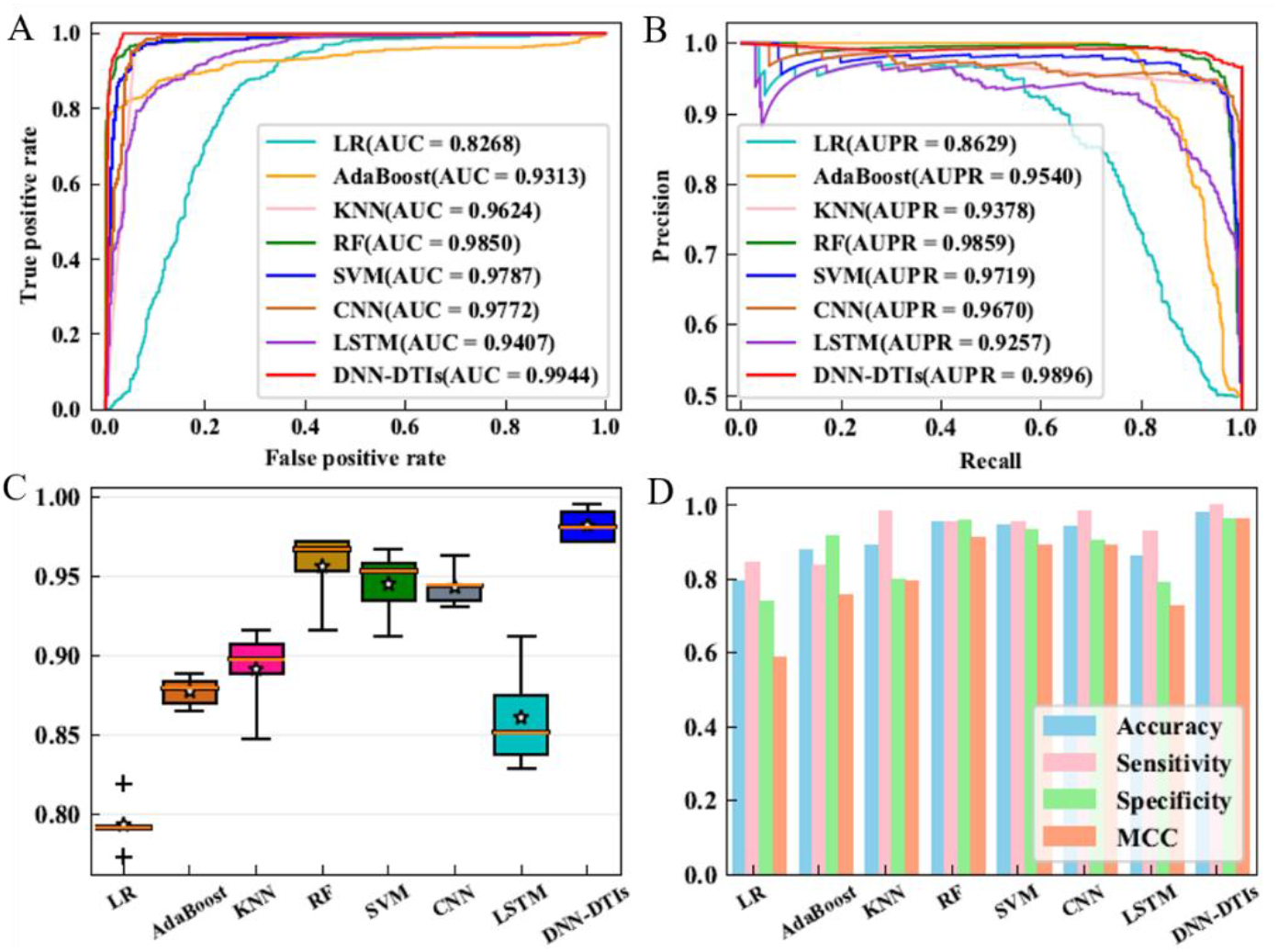
ROC curves (A), PR curves (B), box plots (C) and histograms (D) of LR, AdaBoost, KNN, RF, SVM, CNN, LSTM and DNN-DTIs on NR dataset.

This paper obtained the results of DNN and LR, AdaBoost, KNN, RF, SVM, CNN, and LSTM on the dataset constructed by Kuang et al. The ACC, SE, SP, MCC and AUC values are shown in Table S12. ROC curve, PR curve, box plot and histogram results are shown in Fig. S4. From the above analysis, DNN can effectively mine the nonlinear relationship between drug fingerprint information and target information and categories. These classic classifier algorithms cannot deeply dig into the inherent high-level feature information, and the prediction ability is limited. DL can identify and predict DTIs and non-DTIs through hierarchical learning. Compared with CNN and LSTM, DNN perform better model performance for DTIs prediction.

### 3.5. Comparison with existing DTIs prediction methods

Considering the importance of DTIs, a large number of DTIs methods have been presented, including PDTPS of Meng et al. [65], PSSM+ DVM of Li et al. [66], GIP+ RLS of Laarhoven et al. [67], and BLM-NII of Mei et al. [68], NLRMF of Liu et al. [69], SAR of Cao et al. [19], LRF-DTIs of Shi et al. [26] and Kuang et al. [35]. To ensure the objectiveness of the comparison results, we list the evaluation indicators of DNN-DTIs and others on the same gold standard datasets of Enzyme, IC, GPCR, NR and Kuang et al’s dataset (Table 7).

From Table 7, overall, the performance and indicators of DNN-DTIs are the best. For Enzyme, the AUC values of DNN-DTIs are 6.95%, 2.03%, 1.03%, 1.13%, 4.97%, and 0.01% higher than PSSM+ DVM, GIP+ RLS, BLM–NII, NRLMF, SAR, and LRF-DTIs, respectively (0.9983 VS 0.9288, 0.978, 0.988, 0.987, 0.9486 and 0.9982). The ACC value of DNN-DTIs is 5.62% higher than PSSM+ DVM (98.78% VS 93.16%) and 0.69% higher than LRF-DTIs (98.78% VS 98.09%). For IC, the AUC values of DNN-DTIs are 7.93%, 1.24%, 0.64%, 0.74%, and 5.36% higher than PSSM+ DVM, GIP+ RLS, BLM–NII, NRLMF, and SAR, respectively (0.9964 VS 0.9171, 0.98400, 0.9900, 0.989, 0.9428 and 0.9965). The ACC value of DNN-DTIs is 8.47% better than SAR (98.60% VS 88.91%) and 0.69% higher than LRF-DTIs (98.60% VS 97.32%). For GPCR, The AUC values of DNN-DTIs are 10.84%, 4%, 1%, 2.5%, 10.38%, and 0.22% higher than PSSM+ DVM, GIP+ RLS, BLM–NII, NRLMF, SAR, and LRF-DTIs (0.994 VS 0.8856, 0.954, 0.984, 0.969, 0.8902, 0.9918). The ACC value of DNN-DTIs is 1.28% higher than LRF-DTIs (97.98% VS 95.69%). For NR, The AUC values of DNN-DTIs are 6.44%, 7.24%, 1.34%, 4.44%, 11.22%, and 3.85% higher than PSSM+ DVM, GIP+ RLS, BLM–NII, NRLMF, SAR, and LRF-DTIs (0.9944 VS 0.93, 0.922, 0.981, 0.95, 0.8822 and 0.9559). From the perspective of ACC indicators, the value of DNN-DTIs is 10.46% higher than PDTPS (98.24% VS 87.78%), 6.02% higher than PSSM+ DVM (98.24% VS 92.22%), and 3.36% higher than LRF-DTIs (98.24% VS 94.88%). In terms of AUC, DNN-DTIs are also superior to the method proposed by Kuang et al. (0.9957 VS 0.9350).

As shown in Table 7, the DNN-DTIs achieves the highest performance, comparing with PSSM+ DVM, GIP+ RLS, BLM-NII, NRLMF, SAR, LRF-DTIs and Kuang et al. There are some reasons to illustrate this situation. First, the comprehensive and effective derived fusion featuers can provide good information to predict DTIs. A tree-based feature selection method, XGBoost, can retain better DTIs representation without reducing model complexity. The imbalanced processing algorithm, SMOTE, is employed to synthesize artificial samples, providing optimaized and balanced feaure vectors. In particular, the layer-by-layer learning ability of DNN can effectively mine the non-linear relationship between drug-target interactions information and labels, extracting important and effective high-level feature information.

### 3.6. Prediction of drug-target interactions network

Predicting the DTIs network can contribute to the study of topological property and the identification of new DTIs. DTIs network analysis can visually illustrate the potential biological significance. We use drugs and targets as nodes of the graph. The edges in the graph indicate interactions. This paper uses Network1 and Network2 to demonstrate and verify the DNN-DTIs. First, the raw features from PseAAC, PsePSSM, CTD, CT, NMBroto, structure and substructure fingerprint are concatenated. Then, the optimiazed features are obtained through XGBoost and SMOTE. Finally, DNN-DTIs using NR dataset is constructed to predict Network1 (Fig. 7). The prediction score of each DTI in Network1 can be found in Table S13. As shown in Fig. 8, Network2 is composed of three subnetworks, where Enzyme is represented in orange, IC is represented in green, and GPCR is represented in blue. The gold standard dataset of Enzyme, IC, and GPCR are used as training sets to predict the three subnetworks. The training set eliminates the drug-target interactions pairs that appear in Network2, and the obtained prediction results can be find in Fig. 8.

**Fig. 7.**
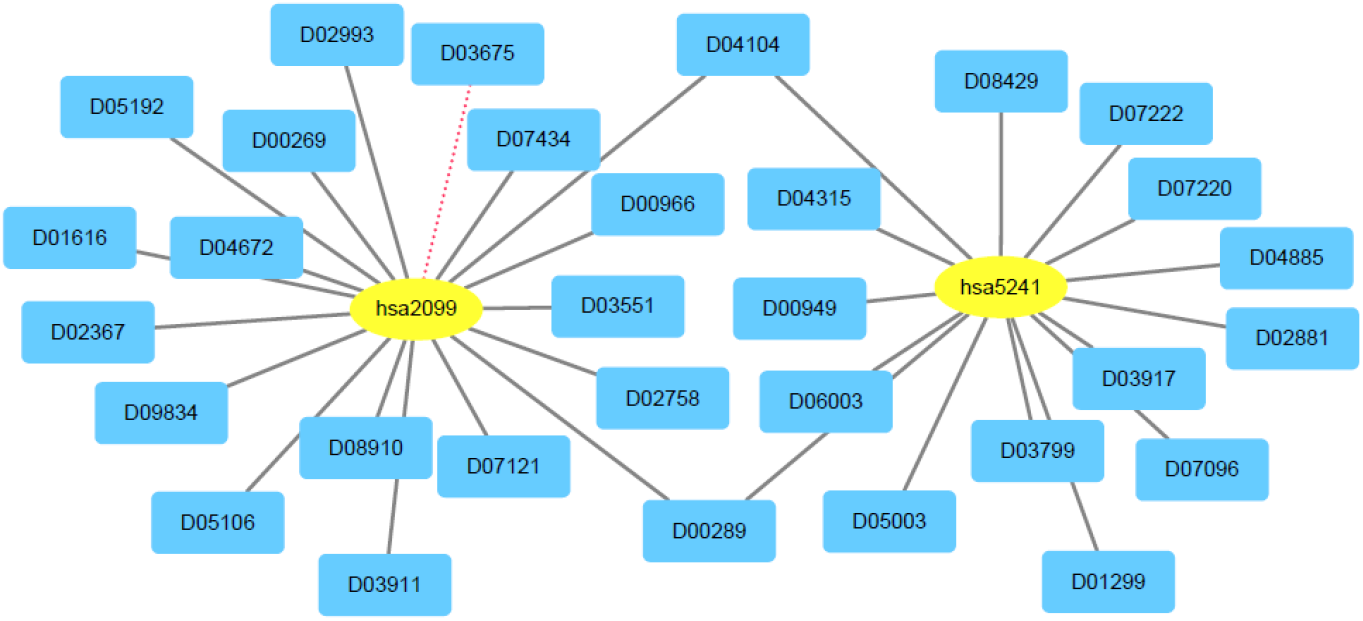
The prediction results of the drug-target interactions Network1. The circular box represents the target, and the rectangular box represents the drug. The solid black edge in the figure indicates that the DTI is predicted successfully, while the red dashed edge indicates that the DTI is not successfully predicted. It can be seen that for 33 pairs of DTIs, only 1 pair failed to be predicted.

**Fig. 8.**
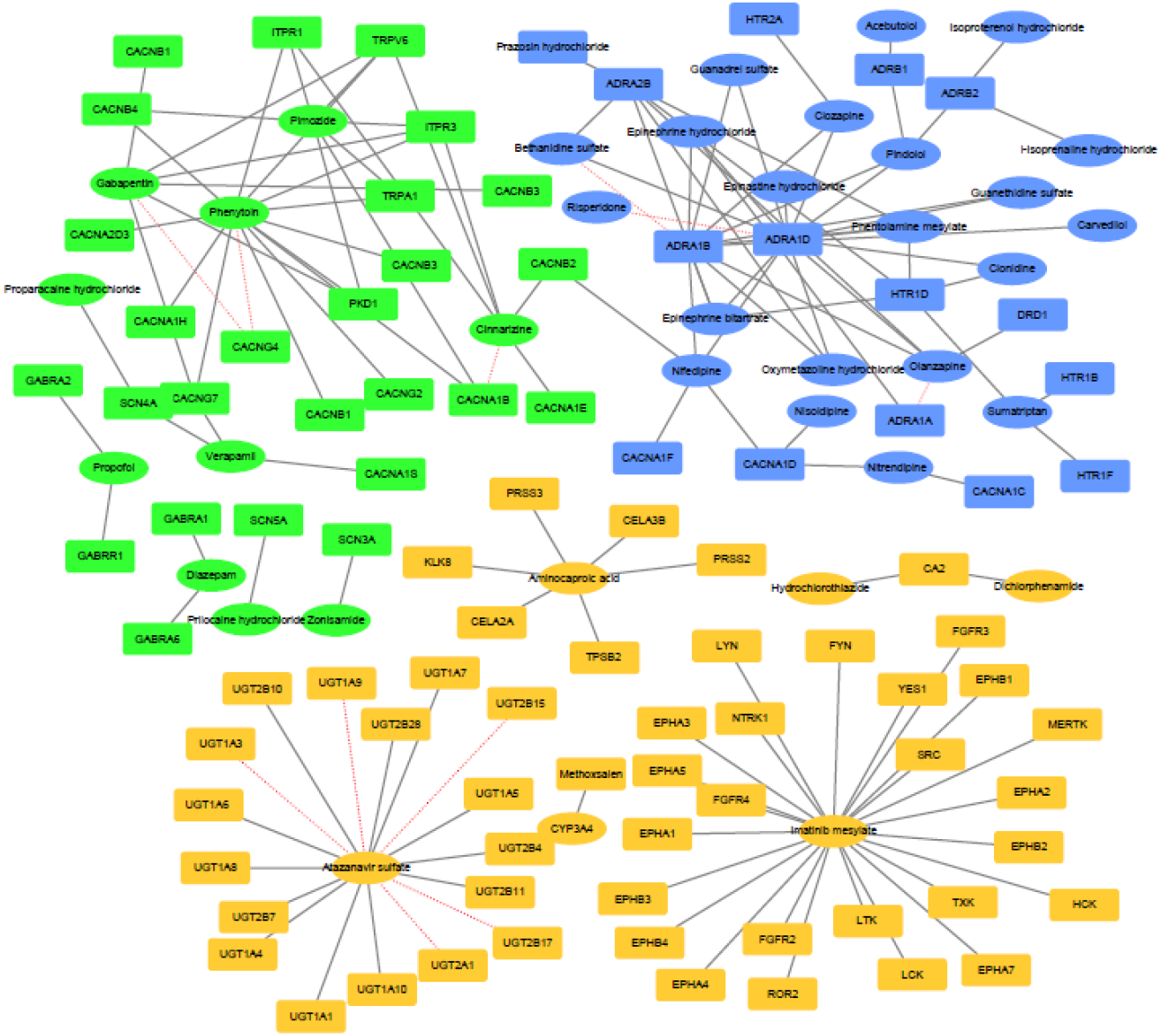
Prediction results of drug-target interactions Network2. The circular box represents the drug, the rectangular box represents the target, the orange part represents subnetwork Enzyme, the green part represents subnetwork IC, and the blue part represents subnetwork GPCR. The solid black line represents the DTI is successfully predicted, and the red dashed edge indicates that the DTI is not successfully predicted. It can be seen that for 150 pairs of DTIs, only 11 pairs of DTIs are not predicted successfully.

Fig. 7 indicates that the trained DNN-DTIs model can effectively predict the DTIs in Network1, and only one pair of DTI D03675-hsa2099 is not predicted successfully (The accuracy is 96.97% (32/33)). While the accuracy of Shi et al. [26] is 85.29% (29/33), DNN-DTIs achieve better performance than LRF-DTIs. The drug name, target name and prediction scores of Network1 are shown in Table S13. From Fig. 8, for Enzyme, UGT2A1-Atazanavir sulfate, UGT1A9-Atazanavir sulfate, UGT1A3-Atazanavir sulfate, UGT2B17-Atazanavir sulfate and UGT2B15-Atazanavir sulfate are not predicted successfully (see orange subnetwork in Fig. 8). For GPCR, ADRA1D-Risperidone, ADRA1B-Bethanidine sulfate and ADRA1A-Olanzapine are not predicted successfully (see blue subnetwork in Fig. 8). Three pairs of drug-target interactions in the IC dataset, CACNG4-Gabapentin, CACNG4-Phenytoin, and CACNA1B-Cinnarizine are not predicted successfully (see the green subnetwork in Fig. 8).

Studies have shown that the combination of the drug Etonogestrel with the receptor can reduce the secretion of luteinizing hormone and can be used to make female contraceptives [70]. The DNN-DTIs prediction method can successfully predict the drug-target interaction D04104-hsa2099 (Etonogestrel-Estrogen receptor). It can be seen that Pimozide in IC dataset is a core drug, and the interaction between Pimozide with PKD1 can be successfully predicted. PKD1 is a gene closely related to polycystic kidney disease. The drug Aglepristone is a progesterone receptor antagonist [71], which can treat various progesterone dependent physiological or pathological states, and has certain effects on diabetes mellitus. Alfatradiol can treat female hair loss [72], obtaining a suitable target can help treat hair loss and diabetes. DNN-DTIs can successfully predict D07096-hsa5241 (Aglepristone-Progesterone receptor) and D07121-hsa2099 (Alfatradiol-Estrogen receptor). According to Enzyme subnetwork, Imatinib mesylate is a small molecule kinase inhibitor which is used to treat cancer-related diseases [73]. DNN-DTIs can predict the interaction between Imatinib mesylate and surrounding targets. The tazanavir sulfate-UGT1A10 interaction is successfully predicted. When tazanavir sulfate binds to the receptor, it can play a therapeutic effect on inflammation and insulin resistance [74]. Pindolol is a 5-HT1A receptor antagonist, which can be used for the design of antidepressants after binding to receptor targets [75]. DNN-DTIs can effectively predict the Pindolol-ADRB1 interaction (the prediction score is 0.9956). Studies have shown that voltage-gated sodium channel is Phenytoin’s main target for anti-arrhythmic drug, and has been used in the treatment of many diseases [76]. DNN-DTIs can successfully predict the Phenytoin-TRPA1 interaction. Therefore, predicting the interactions on Network1 and Network2 can provide new ideas and ways for drug design, drug discovery, and human disease prevention.

## 4. Conclusion

The research and analysis of DTis plays an important role in drug repositioning, drug design and etc. This paper proposes a novel DTIs prediction pipeline called DNN-DTIs based on deep learning. First, we drive fusion vectors containg sequence-based, structure-based, fingerprint-based and evoulution-based information via PseAAC, PsePSSM, CTD, CT, NMBroto, structure feature and substructure fingerprint. DNN-DTIs can preserve effective subset, eliminating redundant features through the tree-based structure and boosting algorithm. Compared with IG, GINI, MRMD, LASSO and EN, XGBoost is outstanding to analyze feature importance. Finally, the optimized features are inputted into DNN to build DNN-DTIs model, where high-level feature and non-linear relationship can be mined through layer-by-layer learning, providing insight and biological significance from raw DTIs information. The results on training set and testing set indicate that DNN-DTIs is good and effective. Finally, DNN-DTIs demonstrates the validation for drug design when it predicts DTIs on Network1 and Network2 successfully. The graph convolutional neural network (GCNN) can mine high-level network topology feature information. Combining GCNN with residual neural networks, capsule neural networks or other popular DL methods to predict drug-target interactions is our next research direction.

## Acknowledgments

This work was supported by the National Nature Science Foundation of China (No. 61863010), the Key Research and Development Program of Shandong Province of China (No. 2019GGX101001), and the Natural Science Foundation of Shandong Province of China (No. ZR2018MC007).

