## Supplementary Tables, Supplementary Figures for "DNN-DTIs: improved drug-target interactions prediction using XGBoost feature selection and deep neural network"

**Table S1**

The numbers of drugs, targets and interactions of five benchmark datasets.

| Datasets | Drugs | Targets | Interactions |
| --- | --- | --- | --- |
| Enzyme | 445 | 664 | 2926 |
| IC | 210 | 204 | 1476 |
| GPCR | 223 | 95 | 635 |
| NR | 54 | 26 | 90 |
| Kuang | 809 | 786 | 3681 |

**Table S2**

Amino acid physicochemical attributes and the division of the amino acids into three groups according to each attribute.

| Attribute | Division | | |
| --- | --- | --- | --- |
| Hydrophobicity_PRAM900101 | Polar: RKEDQN | Neutral: GASTPHY | Hydrophobicity:  CLVIMFW |
| Hydrophobicity_ARGP820101 | Polar: QSTNGDE | Neutral: RAHCKMV | Hydrophobicity: LYPFIW |
| Hydrophobicity_ZIMJ680101 | Polar: QNGSWTDER A | Neutral: HMCKV | Hydrophobicity: LPFYI |
| Hydrophobicity_PONP930101 | Polar: KPDESNQT | Neutral: GRHA | Hydrophobicity: YMFWLCVI |
| Hydrophobicity_CASG920101 | Polar: KDEQPSRNTG | Neutral: AHYMLV | Hydrophobicity: FIWC |
| Hydrophobicity_ENGD860101 | Polar:  RDKENQHYP | Neutral :SGTAW | Hydrophobicity: CVLIMF |
| Hydrophobicity_FASG890101 | Polar: KERSQD | Neutral: NTPG | Hydrophobicity: AYHWVMFLIC |
| Normalized van der Waals volume | Volume range: 0-2.78 GASTPD | Volume range: 2.95-94.0 NVEQIL | Volume range: 4.03-8.08 MHKFRYW |
| Polarity | Polarity value: 4.9-6.2 LIFWCMVY | Polarity value: 8.0-9.2 PATGS | Polarity value: 10.4-13.0 HQRKNED |
| Polarizability | Polarizability value: 0-1.08 GASDT | Polarizability value:0.128-120.186 GPNVEQIL | Polarizability value: 0.219-0.409 KMHFRYW |
| Charge | Positive: KR | Neutral: ANCQGHILMFPSTWYV | Negative: DE |
| Secondary structure | Helix: EALMQKRH | Strand: VIYCWFT | Coil: GNPSD |
| Solvent accessibility | Buried: ALFCGIVW | Exposed: PKQEND | Intermediate: MPSTHY |

**Table S3**

The prediction accuracies of drug-target interactions with different
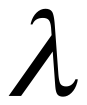
 values on the gold standard dataset.

| Datasets | 5-fold cross-validation (%) | | | | | | | | |
| --- | --- | --- | --- | --- | --- | --- | --- | --- | --- |
|  | 0 | 10 | 20 | 30 | 40 | 50 | 60 | 70 | 80 |
| Enzyme | 93.66 | 95.68 | 95.90 | **96.77** | 95.43 | 87.04 | 87.90 | 85.75 | 84.12 |
| IC | 90.65 | 94.84 | 96.32 | 96.45 | 96.63 | 96.68 | 96.36 | **96.82** | 96.21 |
| GPCR | 93.92 | 95.45 | 95.91 | 95.81 | 96.13 | 96.54 | 96.43 | **96.72** | 96.48 |
| NR | 93.24 | 94.07 | 94.54 | 94.35 | 94.07 | 95.56 | 95.19 | 94.63 | **95.74** |
| Average | 92.87 | 95.01 | 95.67 | **95.85** | 95.57 | 93.96 | 93.97 | 93.48 | 93.14 |

**Table S4**

The prediction accuracies of drug-target interactions with different
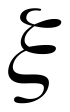
 values on the gold standard dataset.

| Datasets | 5-fold cross-validation (%) | | | | | | | | | | |
| --- | --- | --- | --- | --- | --- | --- | --- | --- | --- | --- | --- |
|  | 0 | 1 | 2 | 3 | 4 | 5 | 6 | 7 | 8 | 9 | 10 |
| Enzyme | **96.34** | 96.15 | 95.74 | 86.91 | 86.57 | 83.26 | 82.05 | 78.70 | 71.31 | 65.00 | 71.98 |
| IC | 94.51 | 95.95 | 96.22 | 96.29 | 96.40 | 95.56 | **97.25** | 96.20 | 96.52 | 96.76 | 96.56 |
| GPCR | 93.16 | 94.57 | 95.22 | 95.37 | 95.22 | 95.39 | **96.10** | 95.46 | 95.55 | 94.54 | 95.20 |
| NR | 90.56 | 93.89 | 93.61 | 94.35 | 93.52 | 94.07 | 93.98 | 93.61 | **95.93** | 95.09 | 93.80 |
| Average | 93.64 | 95.14 | **95.20** | 93.23 | 92.93 | 92.07 | 92.35 | 90.99 | 89.83 | 87.85 | 89.39 |

**Table S5**

The prediction accuracies of drug-target interactions with different
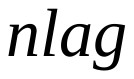
 values on the gold standard dataset.

| Datasets | 5-fold cross-validation (%) | | | | | | | | | |
| --- | --- | --- | --- | --- | --- | --- | --- | --- | --- | --- |
|  | 1 | 2 | 3 | 4 | 5 | 6 | 7 | 8 | 9 | 10 |
| Enzyme | 91.94 | 94.27 | 95.07 | 94.99 | **96.07** | 95.32 | 93.63 | 95.84 | 81.96 | 85.69 |
| IC | 91.89 | 93.82 | 95.44 | 95.84 | 96.71 | 96.58 | 96.48 | 96.64 | **97.09** | 96.74 |
| GPCR | 90.56 | 94.11 | 94.45 | 96.09 | 95.80 | 96.46 | 96.46 | 96.54 | **96.82** | 96.57 |
| NR | 88.33 | 92.41 | 93.24 | 92.31 | 93.70 | 95.09 | 95.28 | 94.07 | **95.83** | 94.63 |
| Average | 90.68 | 93.65 | 94.55 | 94.81 | 95.57 | **95.86** | 95.46 | 95.77 | 92.93 | 93.41 |

Table S6

Prediction results of drug-target interactions on enzyme dataset with different dimensions.

|  | 100 | 200 | 300 | 400 | 500 | 600 | 700 | 800 | 900 | 1000 |
| --- | --- | --- | --- | --- | --- | --- | --- | --- | --- | --- |
| ACC | 98.56 | 98.82 | 98.89 | 98.43 | 97.67 | 97.66 | 85.84 | 75.18 | 72.72 | 61.58 |
| SE | 98.95 | 99.65 | 99.61 | 99.81 | 99.59 | 99.57 | 79.00 | 67.29 | 68.42 | 67.66 |
| SP | 98.18 | 97.99 | 98.18 | 97.04 | 95.74 | 95.76 | 92.69 | 83.08 | 77.03 | 55.49 |
| MCC | 97.14 | 97.66 | 97.80 | 96.89 | 95.41 | 95.40 | 74.68 | 56.72 | 53.54 | 34.58 |
| AUC | 99.81 | 99.86 | 99.85 | 99.75 | 99.42 | 99.36 | 91.86 | 83.47 | 80.24 | 68.40 |

**Table S7**

Prediction results of drug-target interactions on IC dataset with different dimensions.

|  | 100 | 200 | 300 | 400 | 500 | 600 | 700 | 800 | 900 | 1000 |
| --- | --- | --- | --- | --- | --- | --- | --- | --- | --- | --- |
| ACC | 98.26 | 98.48 | 98.51 | 98.48 | 98.24 | 98.08 | 97.84 | 97.61 | 95.26 | 85.32 |
| SE | 99.68 | 99.55 | 99.72 | 99.53 | 99.51 | 99.40 | 99.63 | 99.22 | 99.04 | 98.90 |
| SP | 96.84 | 97.40 | 97.30 | 97.44 | 96.96 | 96.76 | 96.05 | 95.99 | 91.49 | 71.73 |
| MCC | 96.56 | 96.98 | 97.05 | 96.99 | 96.52 | 96.20 | 95.75 | 95.28 | 90.91 | 74.70 |
| AUC | 99.61 | 99.61 | 99.67 | 99.70 | 99.62 | 99.50 | 99.43 | 99.32 | 97.26 | 90.62 |

**Table S8**

Prediction results of drug-target interactions on GPCR dataset with different dimensions.

|  | 100 | 200 | 300 | 400 | 500 | 600 | 700 | 800 | 900 | 1000 |
| --- | --- | --- | --- | --- | --- | --- | --- | --- | --- | --- |
| ACC | 97.69 | 97.64 | 97.86 | 97.40 | 96.89 | 97.06 | 97.14 | 96.42 | 96.77 | 95.66 |
| SE | 99.37 | 99.24 | 99.87 | 99.61 | 99.42 | 99.76 | 99.42 | 99.13 | 99.32 | 98.56 |
| SP | 96.01 | 96.04 | 95.85 | 95.20 | 94.36 | 94.36 | 94.86 | 93.70 | 94.23 | 92.76 |
| MCC | 95.44 | 95.36 | 95.80 | 94.90 | 93.91 | 94.26 | 94.38 | 93.00 | 93.67 | 91.49 |
| AUC | 99.28 | 99.42 | 99.45 | 99.27 | 99.17 | 99.32 | 98.95 | 98.60 | 98.94 | 98.24 |

**Table S9**

Prediction results of drug-target interactions on NR dataset with different dimensions.

|  | 100 | 200 | 300 | 400 | 500 | 600 | 700 | 800 | 900 | 1000 |
| --- | --- | --- | --- | --- | --- | --- | --- | --- | --- | --- |
| ACC | 97.22 | 97.04 | 97.50 | 95.74 | 96.20 | 96.30 | 95.93 | 94.44 | 95.46 | 95.65 |
| SE | 99.26 | 99.07 | 99.26 | 98.89 | 98.89 | 98.70 | 98.33 | 98.52 | 97.59 | 97.78 |
| SP | 95.19 | 95.00 | 95.74 | 92.59 | 93.52 | 93.89 | 93.52 | 90.37 | 93.33 | 93.52 |
| MCC | 94.54 | 94.20 | 95.10 | 91.72 | 92.57 | 92.78 | 91.96 | 89.24 | 91.07 | 91.42 |
| AUC | 98.87 | 99.31 | 99.56 | 98.98 | 98.31 | 98.85 | 98.28 | 97.54 | 98.63 | 98.04 |

**Table S10**

The number of optimal feature subsets retained though different feature selection methods.

|  | Enzyme | IC | GPCR | NR | Kuang |
| --- | --- | --- | --- | --- | --- |
| IG | 300 | 300 | 300 | 300 | 300 |
| GINI | 300 | 300 | 300 | 300 | 300 |
| MRMD | 300 | 300 | 300 | 300 | 300 |
| LASSO | 125 | 221 | 198 | 267 | 140 |
| EN | 143 | 204 | 176 | 260 | 151 |
| XGBoost | 300 | 300 | 300 | 300 | 300 |

**Table S11**

Comparison results of XGBoost and other feature selection methods on the Kuang dataset.

| Feature selection | ACC (%) | SE (%) | SP (%) | MCC |
| --- | --- | --- | --- | --- |
| IG | 78.91 | 78.98 | 78.83 | 0.5788 |
| GINI | 79.96 | 82.02 | 77.89 | 0.6011 |
| MRMD | 88.49 | 94.40 | 82.58 | 0.7769 |
| LASSO | 96.36 | 97.85 | 94.87 | 0.9281 |
| EN | 84.70 | 80.35 | 89.05 | 0.6982 |
| XGBoost | 98.00 | 98.73 | 97.27 | 0.9602 |

**Table S12**

Results of different classification algorithms in Kuang dataset.

| Methods | ACC (%) | SE (%) | SP (%) | MCC | AUC |
| --- | --- | --- | --- | --- | --- |
| LR | 76.53 | 75.87 | 77.20 | 0.5307 | 0.8425 |
| AdaBoost | 90.66 | 90.03 | 91.30 | 0.8133 | 0.9607 |
| KNN | 90.56 | 99.63 | 81.49 | 0.8248 | 0.9684 |
| RF | 90.31 | 89.52 | 91.11 | 0.8081 | 0.9646 |
| SVM | 95.96 | 95.93 | 95.98 | 0.9199 | 0.9901 |
| CNN | 90.06 | 90.28 | 89.83 | 0.8017 | 0.9584 |
| LSTM | 87.30 | 84.08 | 90.51 | 0.7478 | 0.9363 |
| DNN-DTIs | 98.00 | 98.73 | 97.27 | 0.9602 | 0.9957 |

**Table S13**

Drug-target interaction information and their predicted scores in Network1.

| Drug | Target | Score | Drug | Target | Score |
| --- | --- | --- | --- | --- | --- |
| D02758 | hsa2099 | 0.9724 | D04104 | hsa2099 | 0.9655 |
| D07121 | hsa2099 | 0.9952 | D07096 | hsa5241 | 0.9973 |
| D02993 | hsa2099 | 0.9927 | D02881 | hsa5241 | 0.9731 |
| D00269 | hsa2099 | 0.9585 | D01299 | hsa5241 | 0.9731 |
| D03551 | hsa2099 | 0.7964 | D00289 | hsa5241 | 0.9956 |
| D00289 | hsa2099 | 0.9470 | D03799 | hsa5241 | 0.9907 |
| D03675 | hsa2099 | 0.0528 | D03917 | hsa5241 | 0.9616 |
| D02367 | hsa2099 | 0.9797 | D04104 | hsa5241 | 0.9975 |
| D03911 | hsa2099 | 0.9930 | D04315 | hsa5241 | 0.9713 |
| D08910 | hsa2099 | 0.9228 | D00949 | hsa5241 | 0.9748 |
| D04672 | hsa2099 | 0.9938 | D04885 | hsa5241 | 0.9773 |
| D09834 | hsa2099 | 0.8905 | D05003 | hsa5241 | 0.9849 |
| D05106 | hsa2099 | 0.9853 | D07222 | hsa5241 | 0.9904 |
| D05192 | hsa2099 | 0.9531 | D07220 | hsa5241 | 0.9972 |
| D07434 | hsa2099 | 0.8380 | D08429 | hsa5241 | 0.5656 |
| D01616 | hsa2099 | 0.8548 | D06003 | hsa5241 | 0.5138 |
| D00966 | hsa2099 | 0.9615 |  |  |  |

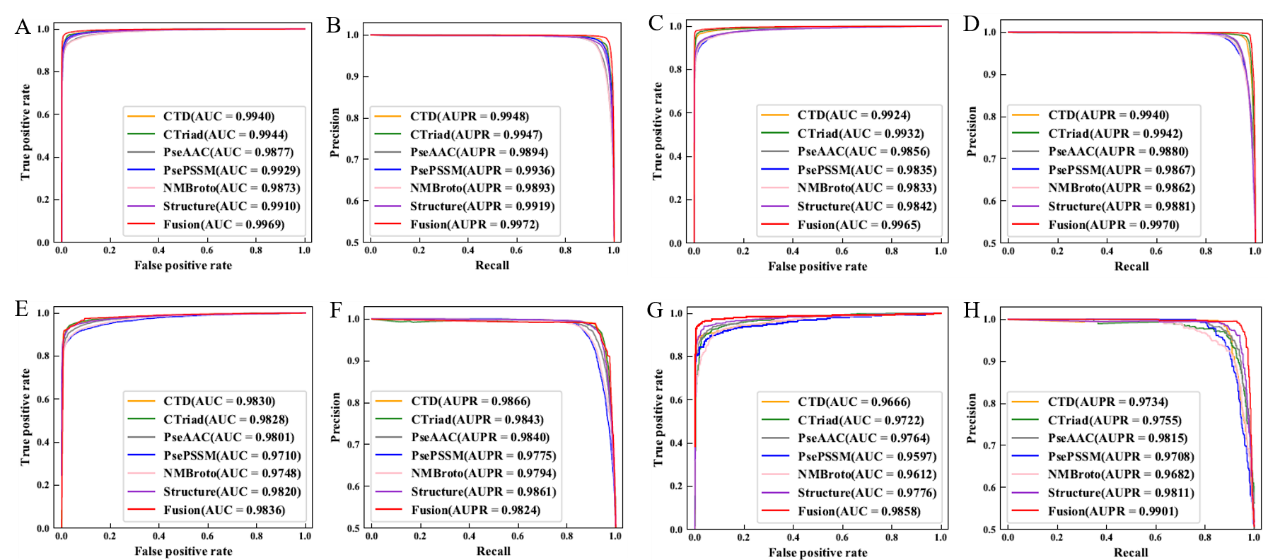

**Fig. S1.** The ROC curve and PR curves of the gold standard dataset with different feature extraction methods. (A-B), (C-D), (E-F) and (G-H) are ROC and PR curves of Enzyme, IC, GPCR and NR, respectively.

Accuracy

| dimension | Enzyme | IC | GPCR | NR |
| --- | --- | --- | --- | --- |
| 100 | 98.56 | 98.26 | 97.69 | 97.22 |
| 200 | 98.82 | 98.48 | 97.64 | 97.04 |
| 300 | 98.89 | 98.51 | 97.86 | 97.5 |
| 400 | 98.43 | 98.48 | 97.4 | 95.74 |
| 500 | 97.67 | 98.24 | 96.89 | 96.2 |
| 600 | 97.66 | 98.08 | 97.06 | 96.3 |
| 700 | 85.84 | 97.84 | 97.14 | 95.93 |
| 800 | 75.18 | 97.61 | 96.42 | 94.44 |
| 900 | 72.72 | 95.26 | 96.77 | 95.46 |
| 1000 | 61.58 | 85.32 | 95.66 | 95.65 |

**Fig. S2.** Histogram of the accuracy of different dimensional models in the gold standard dataset.

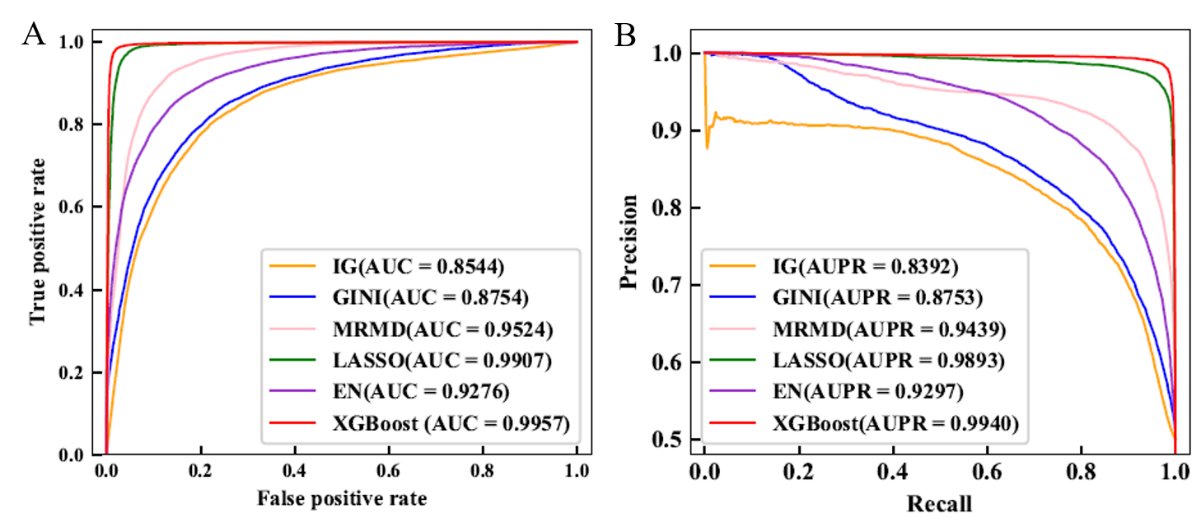

**Fig. S3.** ROC curves (A) and PR curves (B) of Kuang dataset under IG, GINI, MRMD, LASSO, EN and XGBoost feature selection.

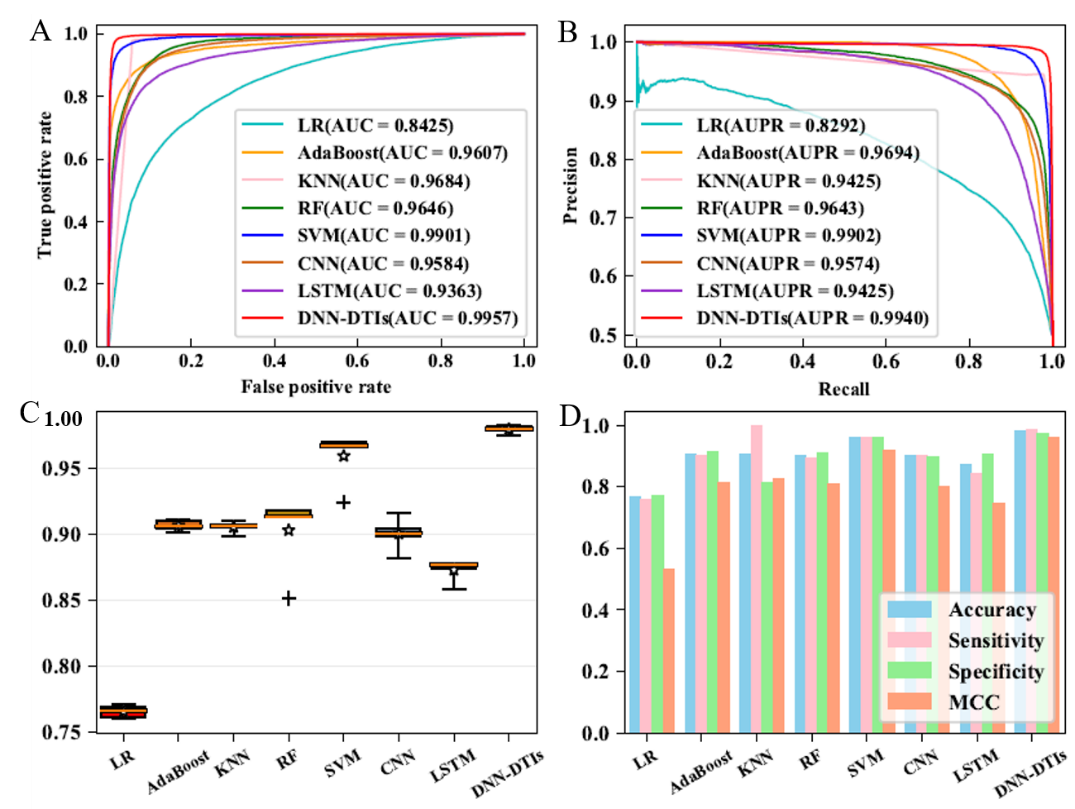

**Fig. S4.** ROC curves (A), PR curves (B), box plots (C) and histograms (D) of LR, AdaBoost, KNN, RF, SVM, CNN, LSTM and DNN-DTIs on Kuang dataset.
